# Mechanism of human tRNA 3’CCA maturation

**DOI:** 10.64898/2026.03.02.709036

**Authors:** Bernhard Kuhle, Luca Krebs, Arjun Bhatta, Sven Dennerlein, Peter Rehling, Hauke S. Hillen

## Abstract

The non-templated addition of the 3’CCA end is the final universal step of tRNA maturation. In humans, 3’CCA addition on nuclear- (nu-tRNA) and mitochondria-encoded tRNAs (mt-tRNA) is catalyzed by a single CCA-adding enzyme, TRNT1, but its precise mechanism remains unknown. Here, we report structures of TRNT1 trapped at various stages during the 3’CCA addition cycle on canonical and non-canonical mt-tRNAs bound to the mitochondrial tRNA maturation platform TRMT10C-SDR5C1. Combined with biochemical data, these structures demonstrate that 3’CCA addition proceeds by a continuous polymerization and translocation mechanism, in which the growing RNA primer remodels the TRNT1 catalytic site to define the specificity of non-templated 3’CCA addition. Moreover, they reveal a relaxed recognition mode that allows TRNT1 to mature both canonical nu-tRNA and non-canonical mt-tRNA substrates. Finally, biochemical analyses of disease-associated TRNT1 variants provide insights into their molecular pathogenesis. Taken together, these results provide a detailed mechanistic picture of human tRNA-3’CCA maturation.

## Introduction

Transfer RNAs (tRNAs) play a central role in cellular gene expression as adaptors between the genetic information stored in messenger RNA (mRNA) and the amino acid sequence of proteins.^1^ They are initially transcribed as precursor transcripts (pre-tRNAs), which are then processed and modified to become functional.^2^ The first steps of tRNA maturation are the removal of leader- and trailer sequences at their 5’- and 3’-ends, respectively, which leave the tRNAs with a single nucleotide overhang at the 3’-end.^2–5^ The tRNA 3’-end is then matured by the non-templated enzymatic addition of C_74_, C_75_, and A_76_,^6,7^ resulting in the universal 3’CCA end that serves as amino acid attachment site and assists peptide bond formation during ribosomal translation.^8–11^ In addition, the 3’CCA acts as a nuclear export signal,^12,13^ serves as a quality control checkpoint for the cellular tRNA pool,^14–17^ and is targeted by nucleases to regulate global translation.^18,19^

The *de-novo* synthesis and repair of 3’CCA ends is catalyzed by a specialized RNA polymerase, the CTP(ATP):tRNA nucleotidyltransferase (CCA-adding enzyme).^6,20^ Its reaction poses unique enzymatic challenges, from the recognition of diverse tRNA substrates to the selection of the correct NTP substrates, switching from CTP to ATP specificity after two rounds of CMP addition and termination once the 3’CCA is complete – all in the absence of a guiding nucleic acid template. CCA-adding enzymes have emerged twice independently, with class I enzymes found in archaea and class II in bacteria and eukaryotes.^21,22^ Previous crystallographic and biochemical analysis of class I enzymes revealed how the protein and tRNA cooperate to select the correct incoming NTP,^7,23–25^ and that 3’-extension proceeds without translocation, instead involving the continuous refolding and scrunching of the growing 3’CCA end in the catalytic cleft.^23,25–27^ By contrast, the mechanism of class II CCA-addition enzymes remains poorly understood. In particular, no high-resolution structures of class II CCA-adding enzymes at intermediate stages of the 3’CCA addition cycle have been reported. As a result, the mechanisms underlying nucleotidyl-transfer, the specificity switch from CTP to ATP, and the structural dynamics of 3’CCA addition remain unclear for class II CCA-adding enzymes.^28^

In human cells, 3’CCA addition is catalyzed by a single class II nucleotidyltransferase named TRNT1.^14,29^ Unlike archaea or bacteria, human cells contain two distinct tRNA pools. While the nuclear genome encodes ∼500 tRNA genes that give rise to ∼300 different tRNAs (nu-tRNAs) used in cytoplasmic protein synthesis,^30,31^ mitochondria additionally contain a minimal set of 22 mitochondrial tRNAs (mt-tRNAs) that are encoded by the mitochondrial genome (mtDNA) and dedicated to mitochondrial translation.^32^ TRNT1 catalyzes 3’CCA addition on both nu- and mt-tRNA pools.^14,29^ All nu-tRNAs share a common ‘canonical’ four-armed cloverleaf-like secondary structure (composed of anticodon-, acceptor-, T- and D-arms) that adopts an L-shaped three-dimensional fold in which the D- and T-loops interact to form the characteristic elbow region.^33–35^ This canonical tRNA structure is highly conserved from prokaryotes to the cytoplasm of human cells and is important for tRNA recognition by archaeal class I and bacterial class II CCA-adding enzymes.^33,36^ By contrast, animal mt-tRNAs underwent a unique process of sequence and structural erosion, which has led to ‘non-canonical’ mt-tRNA structures that lack many of the conserved structural features required for nu-tRNA processing, modification and aminoacylation.^32,37^ Consequently, the enzymes performing these steps are under strong evolutionary pressure to either specialize in recognizing only one of the two tRNA pools or evolve mechanisms to serve tRNAs from both genomes.^38,39^ Recent biochemical and structural studies show that both strategies are employed by the molecular machinery controlling human mitochondrial tRNA processing, maturation, and modification.^38–46^ The 5’-processing of pre-mt-tRNAs is carried out by a dedicated protein-only RNase P composed of the methyltransferase TRMT10C, the dehydrogenase SDR5C1 and the endoribonuclease PRORP.^46,47^ By contrast, 3′-processing of pre-nu-tRNAs and pre-mt-tRNAs is catalyzed by a common RNase Z enzyme named ELAC2.^48,49^ While ELAC2 can cleave canonical tRNAs on its own, it depends on TRMT10C and SDR5C1 to process structurally divergent mt-tRNAs.^39,43^ Previous *in vitro* biochemical data suggest that following 3’-cleavage by ELAC2, the processed mt-tRNA remains bound to TRMT10C-SDR5C1 for subsequent 3’CCA addition by TRNT1,^43^ which is supported by a recent low-resolution cryo-EM structure of TRNT1 bound to TRMT10C-SDR5C1 and mt-tRNA^Ile^.^45^ However, how TRNT1 recognizes its structurally diverse nu- and mt-tRNA substrates and whether it depends on the TRMT10C-SDR5C1 complex to add 3’CCA on non-canonical mt-tRNAs remains unknown.

Here, we combine single-particle cryogenic electron microscopy (cryo-EM) with *in vitro* and cell-based biochemical assays to elucidate the molecular mechanism of human tRNA 3’CCA addition. Cryo-EM structures of TRNT1 stalled at various stages of the reaction cycle with either a canonical (mt-tRNA^Gln^) or non-canonical (mt-tRNA^Tyr^) mt-tRNA substrate bound to TRMT10C-SDR5C1 provide a detailed mechanistic model for 3’CCA maturation. They show that 3’CCA addition by class II enzymes proceeds by a continuous polymerization mode with repeated translocation of TRNT1 along the tRNA, a mechanism that differs from that reported for class I enzymes, and reveal how the growing 3’-end progressively reshapes the enzyme’s active site to determine the specificity of non-templated 3’CCA addition. Moreover, together with mutational and biochemical data, the structures reveal how TRNT1 recognizes structurally divergent tRNAs and demonstrate that TRMT10C-SDR5C1 can promote but are not required for 3’CCA addition on degenerated mt-tRNAs. Finally, biochemical analyses of disease-associated mutations provide insights into the molecular etiology of TRNT1-associated human diseases. Taken together, our results provide a detailed molecular picture of 3’CCA addition on both cytosolic and mitochondrial tRNAs and provide new insights into the function, evolution, and disease-association of the cellular tRNA maturation machinery.

## Results

### Structures of mitochondrial CCA-adding complexes

To determine the molecular basis of tRNA recognition and 3’CCA addition by TRNT1, we reconstituted complexes of TRNT1 with either the canonical mt-tRNA^Gln^ or the structurally eroded, non-canonical mt-tRNA^Tyr^ bound to TRMT10C-SDR5C1 (Figure 1A,B). While complexes with both mt-tRNAs were stable during size exclusion chromatography, complexes with mt-tRNA^Tyr^ disintegrated during grid preparation for cryo-EM. To improve complex stability, the flexible N-terminus of TRMT10C was fused directly to the C-terminus of TRNT1, resulting in a TRNT1-TRMT10C fusion construct that is catalytically active and has improved affinity for canonical and non-canonical mt-tRNAs (Figure S1A-C). To trap TRNT1 in distinct states during the 3’CCA addition cycle, complexes were assembled with mt-tRNA substrates lacking 3’CCA and either with the non-hydrolysable CTP analogue CpCpp (cytidine-5’-[(α,β)-methyleno]triphosphate) (dataset 1), with CTP alone (dataset 2), or with CTP and the non-hydrolysable ATP analog ApCpp (adenosine-5’-[(α,β)-methyleno]triphosphate) (dataset 3).

**Figure 1:**
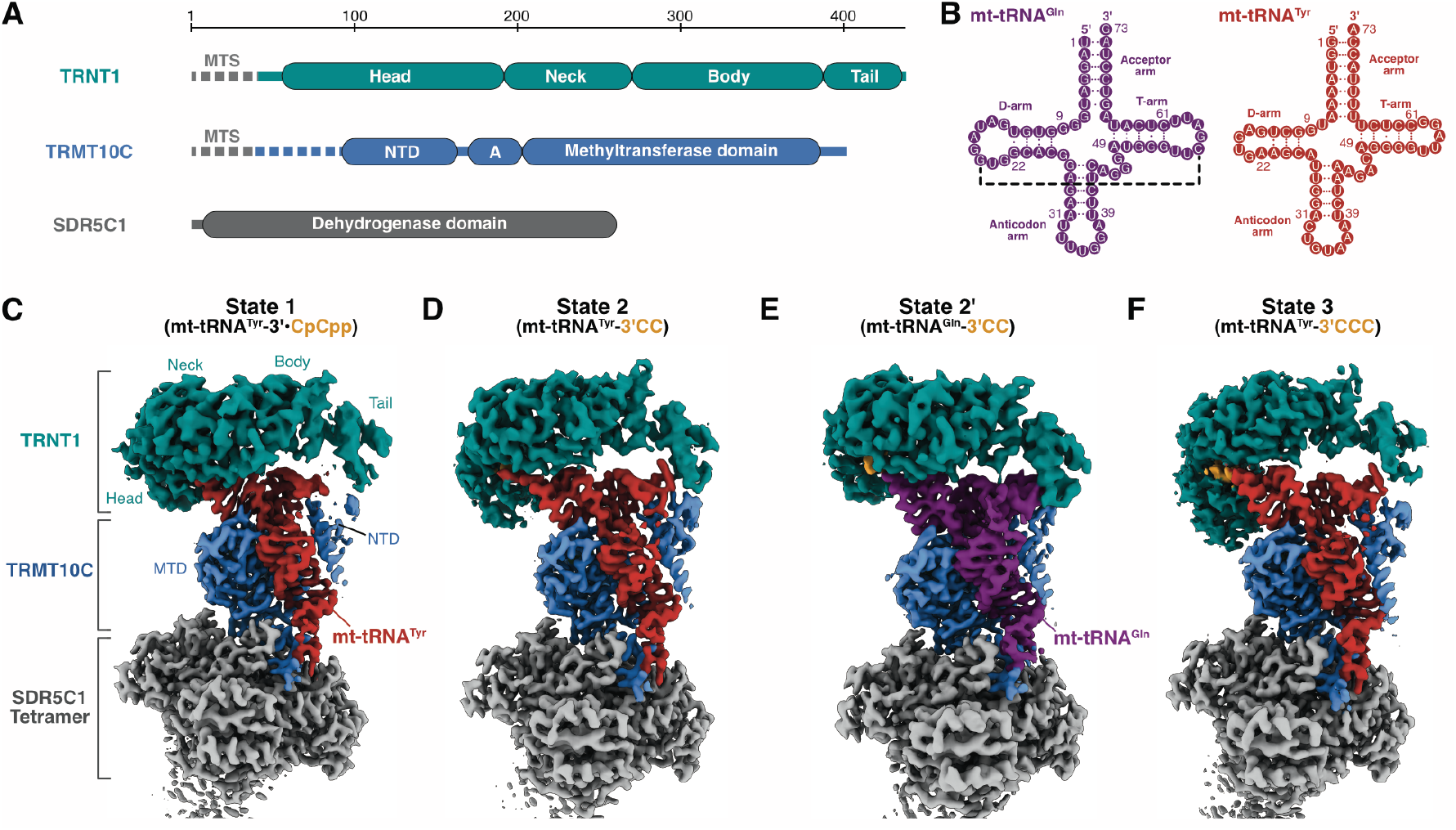
Structures of mitochondrial CCA-adding enzyme complexes. **(A)** Domain representations of TRNT1 (teal), TRMT10C (blue) and SDR5C1 (gray). Unless stated otherwise, this coloring scheme is used throughout the manuscript. Mitochondrial targeting signals (MTS) of TRNT1 and TRMT10C are indicated. The dashed region preceding the NTD of TRMT10C (residues 40-92) is included in the construct used in this study but not resolved in the cryo-EM densities. **(B)** Secondary structures of mitochondrial tRNA^Gln^ (purple) and tRNA^Tyr^ (red) lacking the 3’CCA end. Dashed lines indicate canonical D-loop-T-loop tertiary interactions. The same coloring scheme is used throughout the manuscript. The numbering corresponds to the convention for canonical tRNAs.^87^ **(C)** Cryo-EM map of state 1 of the CCA-adding complex with TRNT1 bound on top of mt-tRNA^Tyr^ and with CpCpp in its active site pocket. **(D)** Cryo-EM map of state 2 of the CCA-adding complex with mt-tRNA^Tyr^-3’CC. The density for the added 3’CC is colored in orange. **(E)** Cryo-EM map of state 2’ of the CCA-adding complex with mt-tRNA^Gln^-3’CC. **(F)** Cryo-EM map of state 3 of the CCA-adding complex with mt-tRNA^Tyr^-3’CCC. See also Figures S1-S7 and Table S1.

Using single-particle cryo-EM, we determined four distinct structures of TRNT1 bound to two distinct mt-tRNAs and TRMT10C-SDR5C1 at global resolutions of 2.6-3.0 Å and local resolutions of 3.0-3.3 Å in the TRNT1 region (Figures 1C-F and S2-6). This allowed us to fit and remodel the structures of TRMT10C, SDR5C1, and TRNT1 based on previously determined mt-RNase Z complex structures^39^ and the AlphaFold model of TRNT1,^50^ resulting in complete structural models of CCA-adding complexes. The four complexes represent different steps of the CCA addition cycle which we denote as states 1-3 according to their sequential occurrence during catalysis. State 1 contains mt-tRNA^Tyr^ ending in 3’A_73_ and CpCpp, state 2 contains mt-tRNA^Tyr^ ending in 3’CC, and state 2’ contains mt-tRNA^Gln^ ending in 3’CC (Figures 2A-D and S7A). In addition to state 2, we observed a second major structural class in dataset 2 (Figure S3), in which mt-tRNA^Tyr^ was extended by three consecutive Cs (-3’CCC, state 3) through the addition of C_76_ instead of A_76_ – a common side reaction of class I and class II CCA-adding enzymes if ATP is not available.^51,52^ Attempts to obtain structures of TRNT1 with mt-tRNA^Tyr^ or mt-tRNA^Gln^ trapped during AMP addition (-3’CC•ApApp) were unsuccessful, as in all cases the majority of picked particles lacked signal for TRNT1, suggesting that it had dissociated either before or during grid preparation. The remaining particles did contain TRNT1, but exclusively in the 3’CC states described above. This suggests that the AMP addition state is inherently transient and that ATP binding may promote dissociation of TRNT1 from the tRNA. Consistent with this, previous reports suggest that the presence of ATP or ApCpp reduces the affinity of TRNT1 for nu-tRNAs by more than an order of magnitude,^53^ and fluorescence anisotropy experiments show that the addition of ApCpp likewise reduces the affinity of TRNT1 for mt-tRNA^Tyr^-3’CC (Figure S1D).

**Figure 2:**
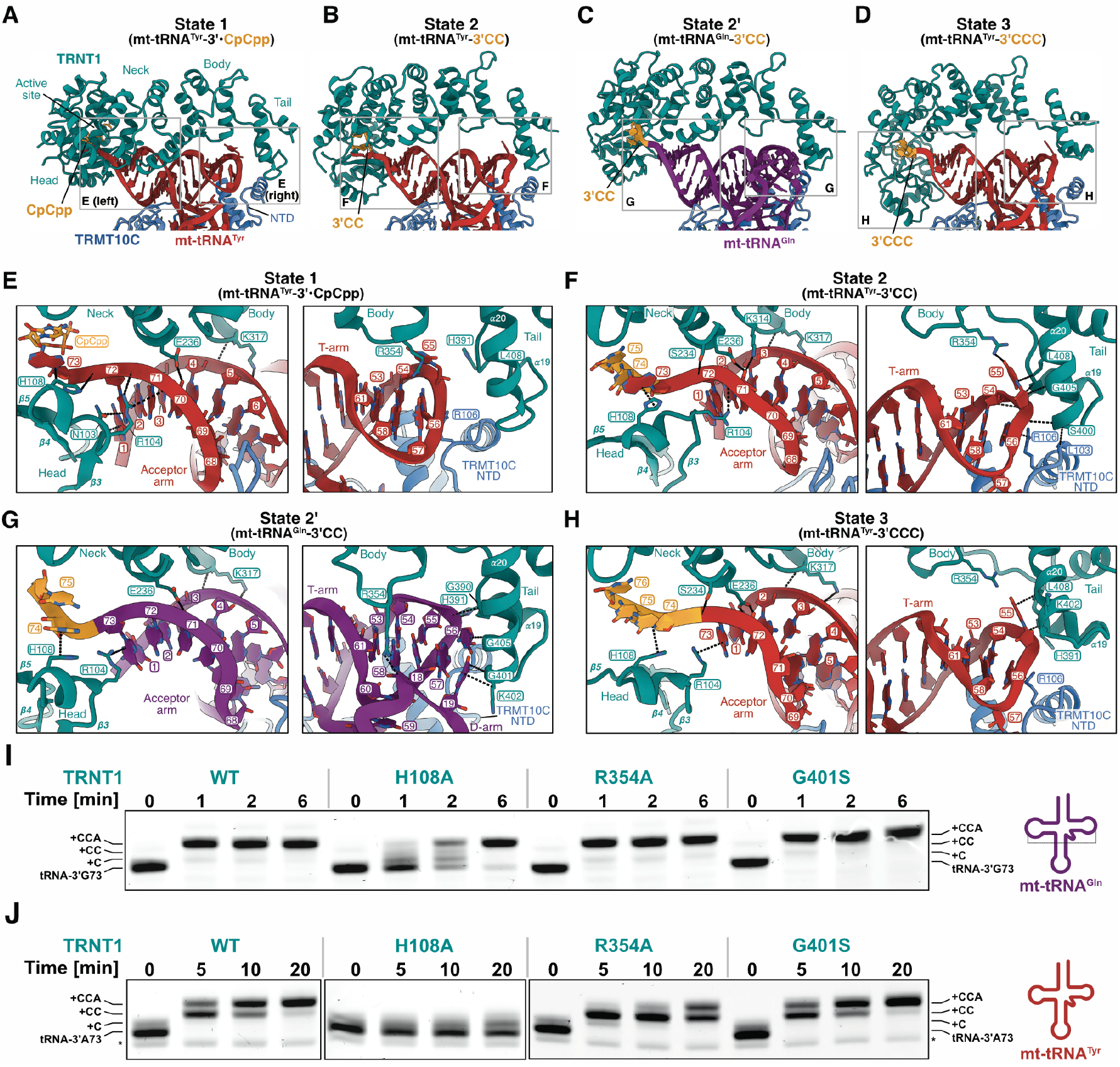
Interactions of TRNT1 with canonical and non-canonical mt-tRNAs. **(A-D)** Structural models of CCA-adding complexes focused on interfaces of TRNT1 with mt-tRNA^Tyr^ and CpCpp in state 1 (A), with mt-tRNA^Tyr^-3’CC in state 2 (B), with mt-tRNA^Gln^-3’CC in state 2’ (C), and with mt-tRNA^Tyr^-3’CCC in state 3 (D). The CpCpp in the active site pocket of state 1 and nucleotides added to the tRNA 3’-end are shown in orange. **(E-H)** Details of the interactions of TRNT1 with the acceptor arm (left) and the elbow region (right) of mt-tRNA^Tyr^ in state 1 (E), mt-tRNA^Tyr^-3’CC in state 2 (F), mt-tRNA^Gln^-3’CC in state 2’ (G), and mt-tRNA^Tyr^-3’CCC in state 3 (H). **(I-J)** *In vitro* activity assays for individual TRNT1 mutants on mt-tRNA^Gln^ (I) and mt-tRNA^Tyr^ (J) in the absence of TRMT10C-SDR5C1. Gels are representatives of 3-4 independent replicates. See also Figures S6 and S7.

All four TRNT1 complexes share the same overall architecture (Figures 1 and S7A), which resembles the mt-RNase P and mt-RNase Z complexes (Figure S7B).^39,40^ The TRMT10C-SDR5C1 platform, composed of an SDR5C1 tetramer to which TRMT10C is anchored through its ‘adapter helix’,^40^ binds each mt-tRNA at its anticodon stem-loop, the elbow region and acceptor stem. As described previously,^45^ TRNT1 binds on top of the tRNA minihelix domain formed by acceptor stem and T-stem-loop, similar to bacterial CC- and A-adding enzymes.^54,55^ Like its bacterial counterparts,^56^ human TRNT1 adopts an elongated overall conformation composed of the N-terminal α/β head domain followed by all-α-helical neck, body and tail domains (Figure S7C).^17,57^ The neck, body and tail domains are primarily responsible for interactions with the tRNA. The head and neck domains are wrapped around the 3’-end of the tRNA, together forming the catalytic cleft and active site pocket (Figure 2).

### Recognition of canonical tRNAs by TRNT1

The structure of TRNT1 bound to mt-tRNA^Gln^-3’CC and TRMT10C-SDR5C1 (state 2’) shows how the CCA-adding enzyme interacts with canonical tRNA substrates (Figure 2C,G). The mt-tRNA^Gln^ adopts a characteristic L-shaped fold, with the canonical tRNA elbow structure formed by tertiary interactions between the D- and T-loops. TRNT1 clamps the acceptor arm–T-arm ‘minihelix’ of mt-tRNA^Gln^ between its N-terminal head and C-terminal tail domains. Compared to the TRNT1 apo structure,^17,57^ the head domain is rotated by ∼5° toward the tRNA, thereby closing the catalytic cleft (Figure S7E). Simultaneously, the tail domain is rotated inwards by ∼40° to interact with the tRNA elbow through Gly_390_ and His_391_ in α_19_, which interact with the sugar-phosphate backbone at C_56_ at the tip of the T-loop, and with Gly_401_, Gly_405_ and Lys_402_ at the N-terminus of α_20_, which stack against the extended aromatic ring system of the conserved G_19_:C_56_ tertiary base-pair (Figure 2C,G). In addition, the side chain of Arg_354_ in the body domain is inserted between the D- and T-loop to interact with A_58_ and G_18_. However, mutational analysis suggests that neither Gly_401_ or Arg_354_ are critical for 3’CCA addition on mt-tRNA^Gln^ (Figure 2I). Notably, hydrogen-bonds by TRNT1 Arg_394_ to TRMT10C Leu_103_ and/or Leu_104_ appear to be the only direct protein-protein interactions between TRNT1 and the TRMT10C N-terminal domain (NTD). Opposite to the elbow region, the acceptor stem enters the active site cavity formed by the head and neck domains (Figure 2C,G). The TRNT1 neck and body domains pack against the acceptor stem with the side chain and backbone of Lys_317_ interacting with the phosphates of A_5_ and G_4_, respectively, and Glu_236_ forming a hydrogen-bond with the 2’-OH of U_71_ . G_73_, the discriminator base of mt-tRNA^Gln^, is stacked between the first base pair of the acceptor stem (U_1_:A_72_) and Arg_104_ in the head domain, while the 3’-end, which is extended by C_74_ and C_75_, is placed within the catalytic cleft. Taken together, the structure with mt-tRNA^Gln^-3’CC shows that TRNT1 recognizes canonical tRNAs through direct interactions with the conserved elbow region and the acceptor stem to position the 3’-end in its active site.

### Recognition of a non-canonical tRNA by TRNT1

The structures of TRNT1 bound to mt-tRNA^Tyr^ reveal how the enzyme recognizes a non-canonical mt-tRNA. To facilitate comparison with the mt-tRNA^Gln^-3’CC complex described above, we here focus on the complex with mt-tRNA^Tyr^-3’CC (state 2). Like mt-tRNA^Gln^, the TRMT10C-SDR5C1 bound mt-tRNA^Tyr^ adopts an L-shaped overall fold, with TRNT1 clamping the minihelix domain between its head and tail domains (Figure 2B,F). In contrast to canonical tRNAs, mt-tRNA^Tyr^ lacks all stable D-loop-T-loop interactions, including the G_19_:C_56_ base pair. The TRNT1 tail domain instead interacts exclusively with the tRNA’s T-loop, which is substantially remodeled compared to mt-tRNA^Tyr^ in the mt-RNase P complex (Figure S8A). While α_19_ binds the sugar-phosphate backbone around U_55_ similar to C_56_ in mt-tRNA^Gln^, the U_55_ base is flipped out from the T-loop of mt-tRNA^Tyr^ and stacks against a hydrophobic surface formed by Gly_405_ and Leu_408_ in α_20_ (Figure 2F). In addition, the flipped-out U_55_ is stabilized by a hydrogen-bond with Arg_354_ in the body domain. Mutation of Arg_354_ to alanine impairs 3’CCA addition on mt-tRNA^Tyr^ (Figure 2J), while it has no effect on mt-tRNA^Gln^ (Figure 2I), suggesting that its interaction is particularly important for the recognition of non-canonical tRNAs. The TRMT10C NTD is rearranged relative to the mt-tRNA^Gln^ complex and moves closer to the TRNT1 tail domain, allowing formation of an alternative hydrogen-bond between TRNT1 Gly_401_ and TRMT10C Leu_104_. At the other end of the minihelix domain, the neck and body domains of TRNT1 pack against the acceptor stem with Lys_317_ contacting the phosphates of A_4_ and A_5_, and Glu_236_ and Lys_314_ forming hydrogen-bonds with C_71_, similar to the situation observed for mt-tRNA^Gln^-3’CC (Figure 2F,G). However, in contrast to mt-tRNA^Gln^-3’CC, in which G_73_ lies outside of the catalytic cleft, A_73_ of mt-tRNA^Tyr^-3’CC is flipped ∼90° into the catalytic cleft and forms a continuous stack with C_74_ and C_75_ in the active site pocket (Figure 2F). In summary, our structural data demonstrate that TRNT1 uses distinct strategies to recognize the divergent elbow regions of canonical and non-canonical mt-tRNAs, while using identical interfaces with the acceptor stems to position the 3’-end for insertion into the catalytic cleft.

### Sequential active site reorganization during catalysis

The structures of TRNT1 bound to mt-tRNA substrates provide sequential snapshots during the 3’CCA extension cycle (Figure 3A and Supplemental Movie S1). The structure of TRNT1 bound to mt-tRNA^Tyr^-3’•CpCpp (state 1) reveals the active site arrangement immediately preceding the addition of the first CMP molecule. Compared to the TRNT1 apo structure,^17,57^ the head domain is rotated by ∼5°, which widens the active site binding pocket and moves β_3-5_ toward the tRNA primer (Figure S8B). This allows β_3_ to form several stabilizing ionic and stacking interactions with the first acceptor stem base pair (G_1_:C_72_) (Figures 3A and S8C). The discriminator base (A_73_) is unstacked from the acceptor stem and flipped ∼90° into the ‘priming site’ (P site) in the catalytic cleft, where it is sandwiched between His_108_ from the loop linking β_3_ and β_4_, which we denote as ‘wedge loop’, and the base moiety of the incoming CpCpp molecule (Figure 3A,D). This interaction appears to be critical for stabilizing and positioning the RNA primer, as mutation of His_108_ to alanine dramatically reduces TRNT1 activity on both mt-tRNA^Gln^ and mt-tRNA^Tyr^ (Figure 2I,J). The flipped-in A_73_ forms no base-specific interactions, explaining how the P site can accommodate all four bases in the discriminator position. As in bacterial class II enzymes,^56,58^ the CTP (CpCpp) molecule is tightly bound in a ‘templating site’ (T site) in the active site pocket, where hydrogen-bonds with the conserved Asp_194_ and Arg_197_ residues enable protein-based nucleotide selection (Figure 3B-D). Additional density suggests that two magnesium ions are coordinated between the α-, β-, and γ-phosphates of CpCpp, the three catalytic residues Asp_77_, Asp_79_, Glu_121_, and the 3’-OH of A_73_ (Figures 3B and S6B). The 3’-OH of A_73_ is placed directly next to the α-phosphate of CpCpp, suggesting that the complex represents a state immediately before C_74_ addition (Figure 3B). At this stage, the active site pocket is too small to accommodate an ATP molecule (Figure 3F), suggesting that TRNT1 exploits the size difference between CTP and ATP to prevent premature AMP addition. Notably, Arg_126_ and Arg_150_, which are inserted into the catalytic cleft and interact with C_74_ and C_75_ at later stages of the 3’CCA addition cycle (Figure 3E,G), are oriented away from the catalytic cleft (Figure 3D,F), and the ‘flexible loop’ region (V_129_-Thr_142_), which is implicated in the specificity switch from CTP to ATP,^58,59^ is completely disordered (Figures 3F and S8D).

**Figure 3:**
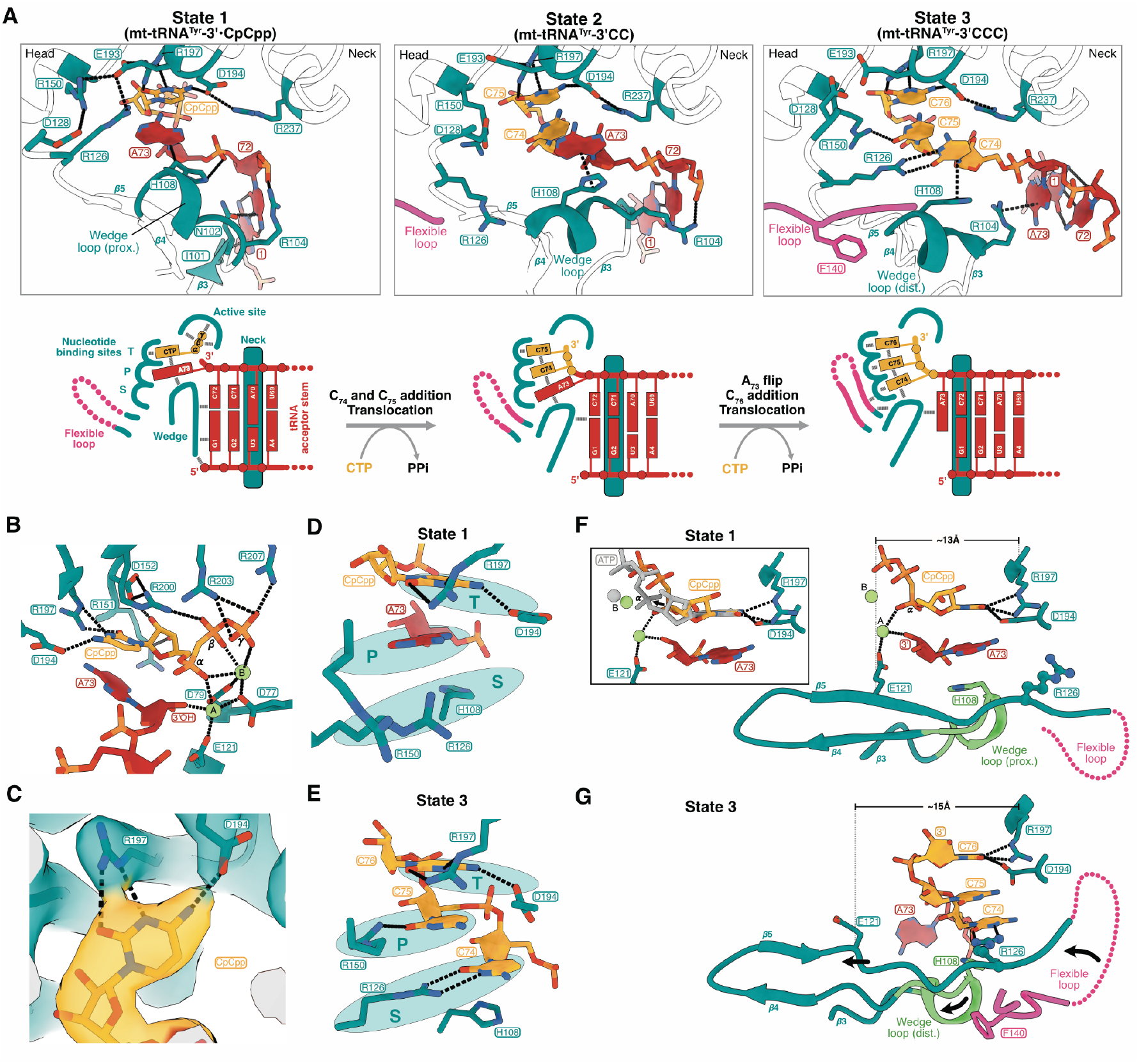
Catalytic cleft organization and dynamics during nucleotidyl-transfer. **(A)** Structures (top) and corresponding schematics (bottom) of the catalytic cleft organization in states 1-3 of TRNT1 bound to mt-tRNA^Tyr^. Structures are superimposed on the neck domain. Black dashed lines indicate interactions of TRNT1 with the acceptor end. **(B)** Active site organization in TRNT1 bound to mt-tRNA^Tyr^-3’ and CpCpp (state 1). Magnesium ions are shown as green spheres. Interactions are indicated by black dashed lines. **(C)** Cryo-EM density and interactions of the incoming CTP (CpCpp) with the protein-based Asp_194_/Arg_197_ template. **(D)** Organization of the templating (T), priming (P), and specificity-determining (S) nucleotide binding sites in the catalytic cleft of TRNT1 in state 1. **(E)** Organization of the three nucleotide-binding sites T, P, and S in the catalytic cleft of TRNT1 in state 3. **(F)** Structural arrangement of the catalytic cleft in state 1. Inset: An ATP molecule (gray sticks) and its bound Mg^2+^ ion (gray sphere) modeled into the active site of state 1 based on the superposition with ATP-bound *B*.*stearothermophilus* CCA-adding enzyme (PDB: 3h39).^56^ The α-phosphate of the ATP is shifted by ∼2 Å away from the tRNA 3’ end relative to the α-phosphate of CpCpp (black arrow). **(G)** Structural arrangement of the catalytic cleft in state 3 (same view as in (F), with structures superimposed on the neck domain). Arrows indicate rearrangements in TRNT1 relative to state 1 that result in and increased size of the active site. See also Figure S8.

The structure of TRNT1 bound to mt-tRNA^Tyr^-3’CC (state 2) represents an intermediate post-catalytic state in which C_74_ and C_75_ have been added to A_73_ (Figure 3A). C_75_ occupies the templating site, while C_74_ resides in the priming site that was previously occupied by A_73_ in state 1. Compared to state 1, the priming site is remodeled by the insertion of Arg_150._ In turn, A_73_ is moved away from the active site pocket into a new binding site, which we denote as ‘specificity-determining site’ (S site), where it still does not form any base-specific interactions (Figure 3A). The catalytic cleft is thus simultaneously occupied by C_75_, C_74_, and A_73_, which together form a continuous stack from the active site pocket to His_108_, which is pushed away from the active site into a ‘distal position’. Although the complex of TRNT1 with mt-tRNA^Gln^-3’CC is likewise stalled in a post-catalytic state with C_74_ and C_75_ added, its active site organization differs from that in the mt-tRNA^Tyr^-3’CC complex. We therefore denote the mt-tRNA^Gln^-3’CC complex as state 2’ (Figure 2G). Most notably, the discriminator base of mt-tRNA^Gln^-3’CC, G_73_, is flipped out of the catalytic cavity and stacks directly against the acceptor stem, C_74_ is moved next to His_108_, and C_75_ partially occupies the priming position. However, as the latter has not fully vacated the templating site, the active site is not yet free to receive an incoming ATP molecule for the final extension step. Taken together, these observations suggest that state 2’ represents an intermediate state following state 2. Overall, the head domain is less well defined in the cryo-EM densities of states 2 and 2’ compared to state 1, suggesting increased flexibility (Figure S8E).

The structure of TRNT1 with mt-tRNA^Tyr^-3’CCC (state 3) provides a model for the complex after addition of three nucleotides to the tRNA 3’-end. Addition of 3’CCC is a common side reaction of class II CCA-adding enzymes if ATP is not available,^51,52^ suggesting that the active site architecture is similar to the general post-catalytic state. The catalytic cavity is occupied by three nucleotides, C_74_, C_75_, and C_76_, that form a continuous stack from the active site pocket to His_108_ in the wedge loop, similar as observed in state 2 (Figure 3A,E). The erroneously added C_76_ occupies the templating site that would otherwise be occupied by A_76_, C_75_ occupies the priming site in the middle and hydrogen-bonds with Arg_150_, and C_74_ occupies the ‘specificity-determining’ (S) site next to His_108_ and forms base-specific interactions with the inserted Arg_126_ (Figure 3E,G). As observed for G_73_ in state 2’, A_73_ is flipped out of the catalytic cavity (Figure 3A), suggesting that its binding capacity is limited to three nucleotides. The translocation of C_74_ into the S site is accompanied by two major structural rearrangements in the head domain of TRNT1 (Figures 3G and S8F): First, the insertion of Arg_126_ and stabilization of the wedge loop in its distal position shift β_3_-β_5_ away from the neck domain, thereby increasing the size of the active site (Figure 3F,G). Second, they allow the ‘flexible loop’ to associate directly with the catalytic cleft entrance in the vicinity to Asp_194_/Arg_197_ (Figure S8D), which is implicated in the specificity switch from CTP to ATP addition.^58,59^

Taken together, the sequential reaction snapshots show that 3’CCA addition is accompanied by the repeated translocation of added nucleotides between three adjacent binding sites within the catalytic cleft. Moreover, they suggest that the translocation of C_74_ into the S site induces a remodeling of the active site that triggers the specificity switch from CTP to the larger ATP.

### TRNT1 rotates and translocates relative to the tRNA during 3’CCA addition

The structural snapshots also provide insights into the global dynamics of 3’CCA addition by TRNT1 (Figures 4 and S9). They reveal that the extension of the 3’-end is accompanied by a continuous, screw-like rotation and translocation of TRNT1 relative to the TRMT10C-SDR5C1-bound tRNA (Figure 4A and Supplemental Movie S2). The start of this motion is represented by the pre-catalytic state 1, in which the neck, body and tail domains of TRNT1 are tilted ∼50° toward the D-loop side of the tRNA. In state 2, the neck, body and tail domains are rotated by ∼30° away from the D-loop and shifted ∼3.8 Å in the direction of the acceptor end. Both movements continue along the same trajectory during transition to state 3. This motion around the tRNA minihelix domain corresponds to a stepwise translocation of TRNT1 by one base pair with each extension at the tRNA’s 3’-end (Figure 4B). The translocation of TRNT1 extends into the tail domain, which moves together with the rest of the protein, causing α_20_ to continuously rearrange relative to the T-loop, sliding over the flipped-out U_55_ (Figure 4C). Additional sub-states of the mt-tRNA^Tyr^-3’CCC complex (Figure S9B) suggest that this trend continues until the tail domain is nearly completely released from the tRNA elbow. Despite the rearrangement of the discriminator base (N_73_) between states 2 and 2’, TRNT1 adopts the same conformation relative to the tRNA in both complexes (Figure 2F,G), suggesting that the observed translocation proceeds similarly for both canonical and non-canonical tRNA substrates. In summary, our data demonstrate that 3’CCA addition by TRNT1 is accompanied by the continuous translocation of the enzyme relative to the tRNA and suggest a mechanism for the release of TRNT1 by translocation-induced dissociation.

**Figure 4:**
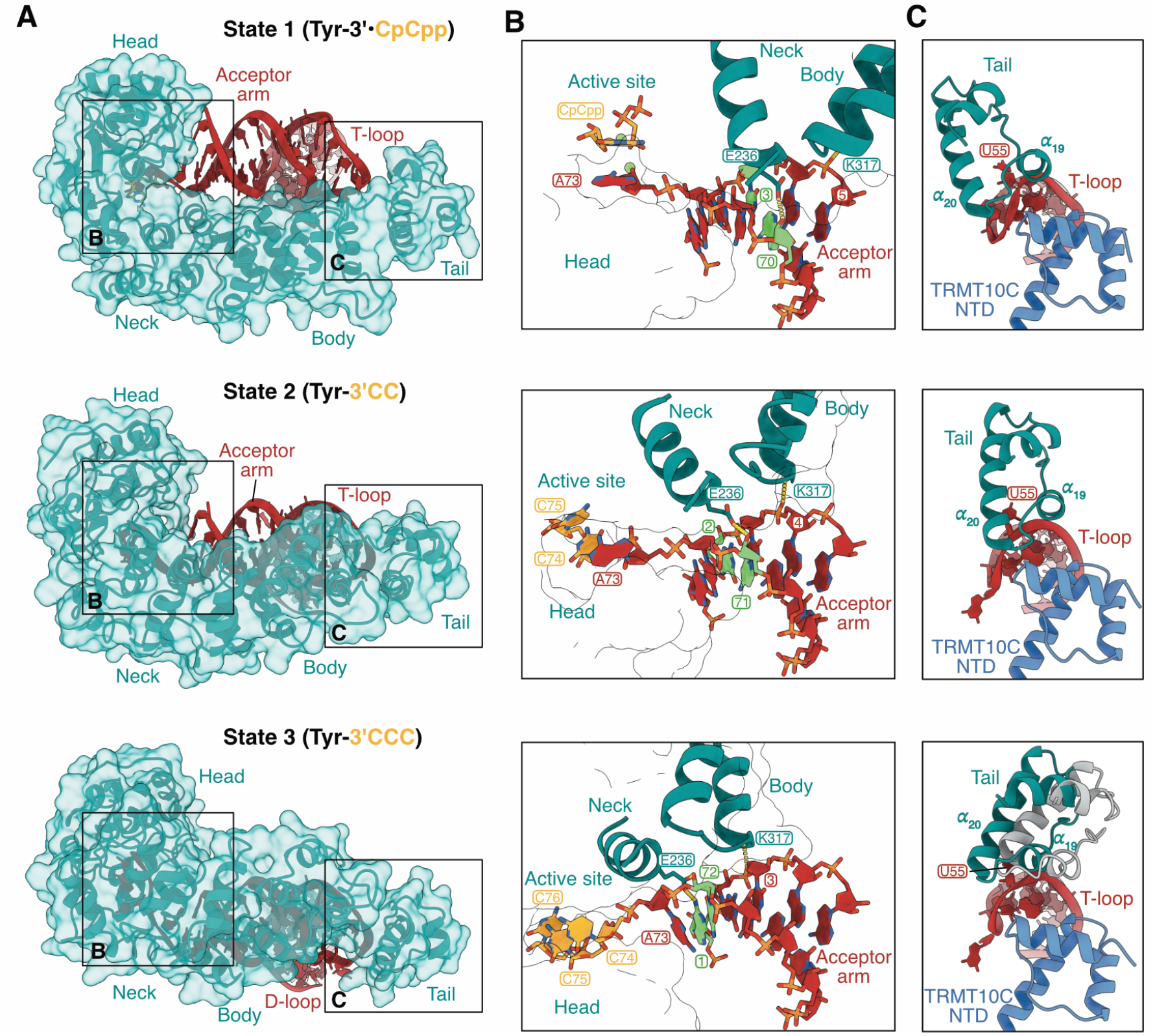
Translocation of TRNT1 during 3’-extension. **(A)** Global movement of TRNT1 relative to mt-tRNA^Tyr^ between state 1 (top), state 2 (middle), and state 3 (bottom). Structures are superimposed on the tRNA. **(B)** Details of the interactions between the neck and body domains of TRNT1 and the acceptor arm of mt-tRNA^Tyr^ in states 1 (top), 2 (middle), and 3 (bottom). In each state the base-pair contacted by Glu_236_ is colored in green. Structures are superimposed on the tRNA. **(C)** Comparison of the interactions of the TRNT1 tail domain with the T-loop of mt-tRNA^Tyr^ in states 1 (top), 2 (middle), and 3 (bottom). Structures are superimposed on the tRNA. In state 3, an additional sub-state is shown in white. See also Figure S9.

### TRMT10C-SDR5C1 is not essential for 3’CCA addition on degenerated mt-tRNAs

In mitochondrial RNase P and RNase Z complexes, the catalytic subunits PRORP and ELAC2 form stable protein-protein interactions with the TRMT10C NTD that promote their catalytic activities on pre-mt-tRNAs.^39,40,43^ No comparable interactions are formed between TRNT1 and TRMT10C (Figure 2), and none of the potential direct interactions are stable throughout the CCA-addition cycle, as the TRNT1 tail domain continuously reorients relative to the TRMT10C NTD (Figure 4C). This suggests that TRNT1 does not require TRMT10C and SDR5C1 for the maturation of structurally divergent mt-tRNAs. To confirm this, we assayed 3’CCA addition by TRNT1 on six mt-tRNAs representing the entire spectrum of structural idiosyncrasies, from an enlarged D-loop in mt-tRNA^Leu(UAA)^ to the smallest D-loop in mt-tRNA^Lys^, the smallest T-loop in mt-tRNA^Thr^ or the complete absence of the D-arm and an extended T-arm in mt-tRNA^Ser(GCU)^.^42^ The results demonstrate that TRNT1 efficiently catalyzes 3’CCA addition on all six mt-tRNA substrates, regardless of the presence or absence of TRMT10C-SDR5C1 (Figure 5A).

**Figure 5:**
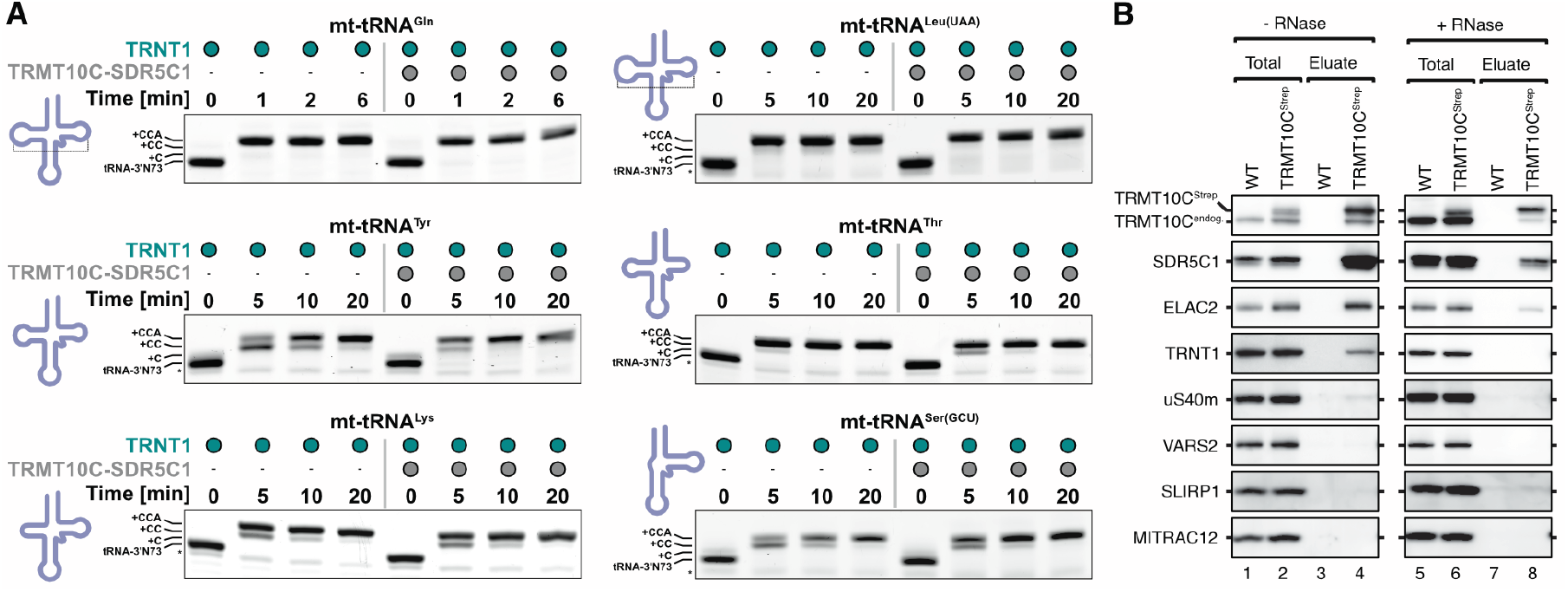
TRNT1 does not require TRMT10C-SDR5C1 for 3’CCA addition. **(A)** *In vitro* activity assays for 3’CCA addition by TRNT1. Gels are representatives of 3-4 independent replicates. **(B)** Western blots of the pull-down of TRMT10C^Strep^ from HEK293T cells. Blots are representatives of two independent replicates.

### TRNT1 binds to TRMT10C-SDR5C1-mt-tRNA in cells

While TRNT1 does not require TRMT10C-SDR5C1, our *in vitro* data show that the presence of TRMT10C-SDR5C1 does not interfere with TRNT1 activity either and can even promote 3’CCA addition on individual degenerated mt-tRNAs such as mt-tRNA^Tyr^ (Figure 5A). This supports the hypothesis that 3’CCA addition occurs while the tRNA is still bound to the TRMT10C-SDR5C1 platform.^43^ To test whether TRNT1 binds TRMT10C-SDR5C1-bound mt-tRNAs *in vivo*, we expressed C-terminally strep-tagged TRMT10C in HEK293T cells at endogenous levels and performed pull-down experiments coupled to western blot analysis (Figure 5B). The results show that TRNT1, like ELAC2, interacts with TRMT10C under cellular conditions. Consistent with our observations that the interaction is almost entirely mediated by tRNA, RNase treatment leads to the loss of TRNT1 binding, whereas ELAC2, which forms more extensive protein-protein interactions with TRMT10C,^39^ is still pulled down (Figure 5B). These results suggest that TRNT1 associates with mt-tRNAs on the TRMT10C-SDR5C1 platform under physiological conditions.

### A structural framework for disease-causing mutations

TRNT1 has a strong association with human disease, with more than 20 homo- and heterozygous mutations in TRNT1 linked to various human pathologies (Figures S10A).^60–62^ To understand how these mutations affect TRNT1 activity, we purified four disease-associated TRNT1 variants containing mutations T110I, D128G, A148V or R190I (Figure 6). Thermal melting analysis shows that T110I and D128G only slightly affect the stability of TRNT1, whereas A148V and R190I reduce its T_m_ by about 7 and 6°C, respectively (Figure 6B). In 3’CCA-adding activity assays with mt-tRNA^Tyr^ and mt-tRNA^Gln^ (Figures 6C,D), A148V showed almost no effect compared to wild-type TRNT1, D128G specifically abolished ATP addition, T110I reduced both CTP and ATP addition, and R190I abolished ATP addition on mt-tRNA^Tyr^ but not on mt-tRNA^Gln^. Overall, the negative effects of TRNT1 mutations appear to be stronger on the non-canonical mt-tRNA^Tyr^ than on canonical mt-tRNA^Gln^, which may explain the dominance of mitochondrial dysfunction in the clinical presentations of TRNT1-associated diseases.^61–63^ Together with our structural data, these results suggest that disease-causing TRNT1 mutations can be subdivided into three main classes (Figures 6 and S10): 1. Mutations like A148V that lie in the structural core of the head and neck domains and reduce cellular TRNT1 activity mainly by local misfolding and/or reduced thermal stability. 2. Mutations like T110I that lie within the catalytic cleft and interfere directly with substrate binding and/or nucleotidyl-transfer without affecting thermal stability. 3. Mutations like D128G and R190I that lie in the interface between the TRNT1 head and neck domains and affect CCA-addition indirectly by altering the structural dynamics of the catalytic cleft (Figures 6 and S10C). Notably, our data show that TRMT10C-SDR5C1 can partially compensate for general defects in 3’CCA adding activity caused by class 1 and 2 mutations, whereas it appears to be unable to restore defects that lead to a specific loss of ATP addition (Figure 6C,D). In summary, our structural data provide a framework to understand the molecular etiology of TRNT1-associated diseases.

**Figure 6:**
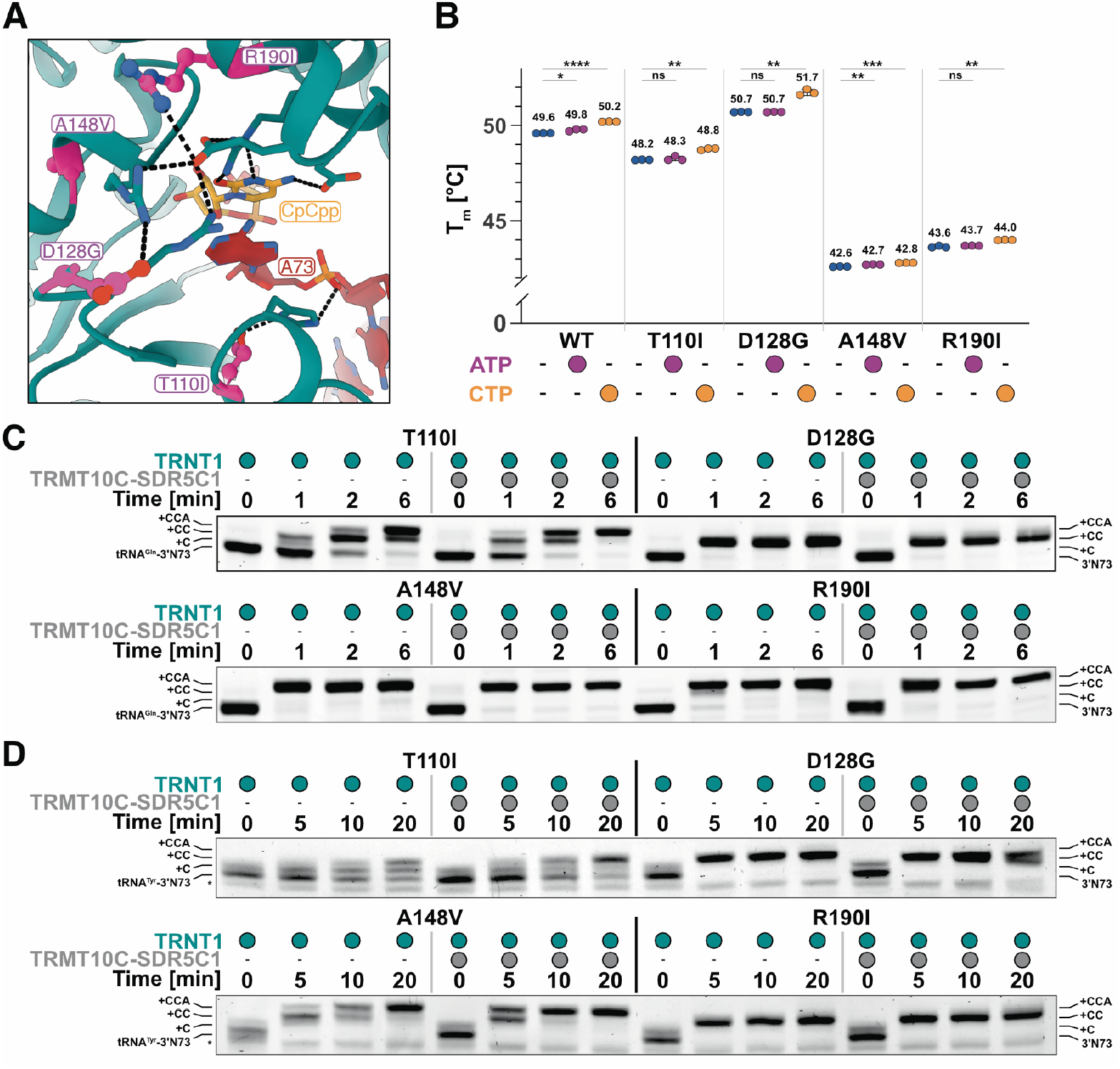
Characterization of disease-causing TRNT1 mutations. **(A)** Disease-causing mutations in the catalytic cleft of TRNT1. Mutation sites are highlighted in pink with wild-type side chains shown in ball-and-stick presentation. **(B)** Thermal melting temperatures (T_m_) of wild-type and mutant TRNT1 variants. Experiments were performed in technical triplicates. Standard deviations from the mean are indicated by error bars. Statistical significance was determined by two-tailed t-test. **(C-D)** *In vitro* CCA-adding assays with mt-tRNA^Gln^ (C) and mt-tRNA^Tyr^ (D) for individual disease-causing TRNT1 mutants in the absence or presence of TRMT10C-SDR5C1. The asterisk marks an RNA band corresponding in length to mt-tRNA^Tyr^ lacking 3’-A_73_. Gels are representatives of two independent replicates. See also Figure S10.

## Discussion

Here we present a detailed structural analysis of the human class II CCA-adding enzyme TRNT1 in complex with structurally divergent mt-tRNA substrates and the mitochondrial tRNA maturation platform TRMT10C-SDR5C1. Together with mutational and biochemical data, these structures provide insight into the mechanism and structural dynamics of nucleotide selection and transfer and demonstrate that 3’CCA-addition is catalyzed by a continuous polymerization mode with the repeated translocation of TRNT1 relative to the tRNA (Supplemental Movies S1 and S2). Moreover, they reveal how TRNT1 recognizes canonical and non-canonical tRNA substrates, providing the molecular basis of both nucleo-cytoplasmic and mitochondrial 3’CCA addition (Figure 7).

**Figure 7:**
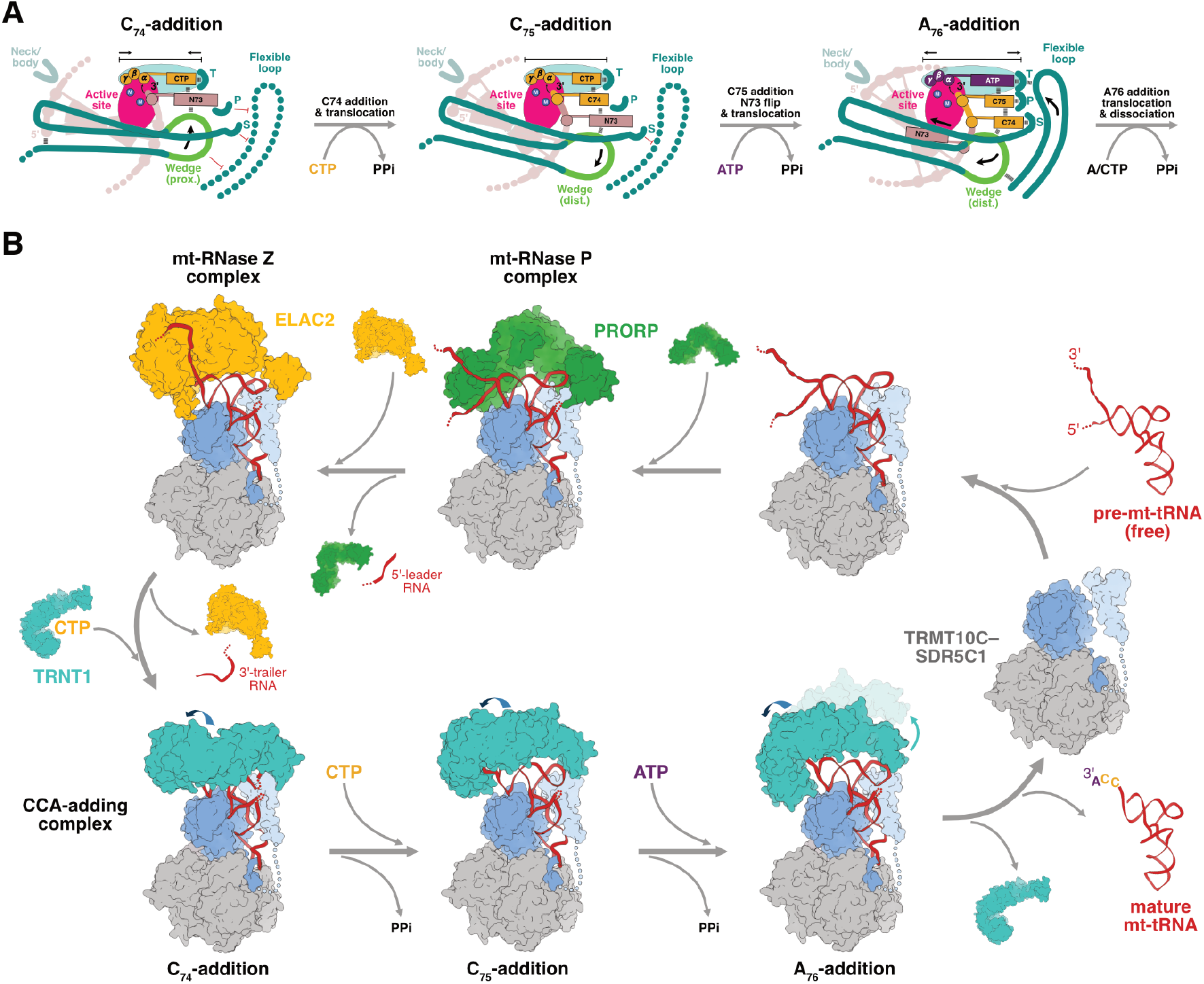
Model of mitochondrial tRNA maturation. **(A)** Model for the structural rearrangements in the catalytic cleft and active site and the transition from CTP to ATP specificity during tRNA 3’CCA addition by TRNT1. **(B)** Model of sequential tRNA maturation in human mitochondria.

The structure of TRNT1 bound with a pre-tRNA and CpCpp (state 1) elucidates the catalytic mechanism of nucleotidyl-transfer by a class II CCA-adding enzyme. Substrate binding to TRNT1 induces the closure of the catalytic cleft and formation of the active site around the flipped-in A_73_ and incoming CTP molecule. A similar closure movement of the head domain was observed in the bacterial class II CC- and archaeal class I CCA-adding enzymes,^7,28^ suggesting a conserved mechanism of substrate-induced active site formation. The active site organization of TRNT1 closely resembles that of structurally homologous palm domains in substrate-bound DNA polymerase β and class I CCA-adding enzymes,^64,65^ supporting a two-cation-dependent catalytic mechanism in which one Mg^2+^ ion activates the 3’-OH of the flipped-in A_73_ for an in-line nucleophilic attack on the α-phosphate and the second Mg^2+^ ion stabilizes the developing negative charge on the leaving group.

Our structures moreover reveal that 3’CCA addition by TRNT1 proceeds by a continuous polymerization and translocation mechanism (Figure 7A and Supplemental Movie S2), akin to template-dependent DNA/RNA polymerases^64,66–68^ but distinct from the refolding and scrunching mechanism observed in class I CCA-adding enzymes.^23–25^ Following the first round of nucleotidyl-transfer, C_74_ translocates twice within the catalytic cleft from the templating (T) site to the priming (P) site and finally to the ‘specificity-determining’ (S) site. The extension of the 3’-end is accompanied by the repeated translocation of the neck and body domains of TRNT1 along the tRNA’s acceptor stem. The wedge loop plays a key role in this process, acting as a counter bearing between the 3’-end inside and the acceptor stem outside of the catalytic cleft, both ensuring the correct placement of the RNA primer in the active site and dynamically adjusting its own position to the number of nucleotides in the catalytic cleft (Figures 2 and 3). Like the alanine mutation of His_108_ in TRNT1 (Figure 2I,J), alanine mutations of the corresponding Phe_85_ and Phe_61_ abolish 3’CCA addition by the *T. maritima* CCA- and *A. aeolicus* A-adding enzymes, respectively,^58,69^ suggesting that the function of the wedge loop is conserved across class II CCA-adding enzymes.

A fundamental unresolved question is how CCA-adding enzymes switch specificity from CTP to ATP after two rounds of CMP incorporation.^7,23,56,58,59,70^ The structure of state 1 shows that a tRNA ending in A_73_ induces an active site conformation that provides a tight, productive fit for an incoming CTP molecule but is too small for an ATP molecule to correctly position its α-phosphate for AMP transfer (Figure 3F). Thus, like class I CCA-adding enzymes,^23^ TRNT1 likely exploits the size difference between CTP and ATP to prevent erroneous AMP incorporation. The structure of state 3 suggests that the translocation of C_74_ into the S site is the key event in the switch from CTP to ATP. C_74_ in the S site induces a concerted structural rearrangement in the TRNT1 head domain that both enlarges the active site pocket and allows association of the flexible loop, which plays a key role in ATP selection,^58,59^ to the catalytic cleft entrance near the templating Asp_194_/Arg_197_ dyad (Figures 3G and SD). This suggests that the specificity switch from CTP to ATP is accomplished by directly coupling the size and shape of the active site pocket to the elongation state of the tRNA 3’-end (Figure 7A). Mutational studies of the *T. maritima* class II CCA-adding enzyme show that Arg_104_, which corresponds to Arg_126_ in the S site of human TRNT1 (Figure 2E), is specifically required only for ATP addition, whereas mutation of Arg_128_ (Arg_150_ in the P site of human TRNT1) abolishes both CTP and ATP addition.^58^ This suggests that the role of the S site to sense the presence of C_74_ and induce the switch from CTP to ATP addition may be conserved in bacterial class II CCA-adding enzymes.

The continuous polymerization mode by TRNT1 also suggests a mechanism for the release of mature tRNA once 3’CCA addition is complete. In contrast to the static interface between class I CCA-adding enzymes and tRNAs,^23,25^ TRNT1 dynamically changes its interactions with the tRNA as the head and neck domains move along the acceptor stem and away from the T-loop, resulting in the gradual loss of protein-RNA interactions (Figures 2 and 4). This suggests that the release of TRNT1 from the mature tRNA results from a translocation-induced dissociation – or ‘unscrewing’ – of the head and tail domains from the tRNA, similar to what has been proposed for the bacterial CC-adding enzyme.^54^ The final trigger for release may be the binding of another incoming NTP into the active site: As the binding capacity of the catalytic cleft is limited to three nucleotides, this would require A_76_ to move into the P site and C_74_ to flip out of the catalytic cleft, thereby further destabilizing the interactions of the head and tail domains with the tRNA’s minihelix domain. This model of translocation-induced dissociation would explain why ATP and ApCpp reduce the affinity of TRNT1 for nu-tRNA-3’CC by more than an order of magnitude and how the release of mature tRNA-3’CCA is promoted by the binding of another ATP to the active site.^52,53^

The structures of TRNT1 bound with different mt-tRNAs also reveal how a single CCA-adding enzyme can interact with the structurally divergent repertoire of human nu- and mt-tRNAs. Bacterial and archaeal CCA-adding enzymes of either class recognize tRNAs by their common overall fold – the uniform length of their acceptor-arm-T-arm minihelix, the consensus sequence and size of the T-loop and their ability to form a canonical elbow structure.^26,36,71,72^ However, most human mt-tRNAs contain highly divergent T-loop sequences and lack a canonical elbow structure.^32,37^ The mitochondrial tRNA processing enzymes PRORP and ELAC2 adapted to this problem by co-opting the auxiliary TRMT10C-SDR5C1 platform into their respective mt-RNase P and mt-RNase Z complexes.^39,40,43,46^ In both cases, TRMT10C-SDR5C1 serve as a platform that stabilizes mt-tRNAs in a common L-shaped fold and provides protein-interfaces as anchor points for PRORP and ELAC2 to compensate for the absence of the canonical elbow.^39,43,47^ TRNT1 does not show the same degree of dependency on TRMT10C-SDR5C1 but catalyzes 3’CCA addition on both free and TRMT10C-SDR5C1-bound mt-tRNAs (Figure 5).^29,43^ Our structures show how TRNT1 interacts with both canonical and non-canonical tRNAs using distinct but overlapping interfaces on its tail domain (Figure 2). The interactions of TRNT1 with the elbow of mt-tRNA^Gln^-3’CC are very similar to those by the *A. aeolicus* CC-adding enzyme with its tRNA substrate,^54^ suggesting a conserved recognition mechanism for canonical tRNAs. By contrast, mt-tRNA^Tyr^ lacks all tertiary D-loop-T-loop interactions and the TRNT1 tail domain instead recognizes the flipped-out U_55_ in a substantially remodeled T-loop. It is important to note, however, that each complex represents only a snapshot during 3’CCA addition (Figure 4), and both the extensive rearrangements of the tail domain as well as the sequence and structural diversity among mt-tRNA variants make it likely that TRNT1 uses more than two interaction modes for canonical and non-canonical tRNAs. Our analysis thus provides another example of how the structural idiosyncrasies and intrinsic flexibility of mt-tRNAs are exploited by nuclear-encoded binding partners.^38,41,42^ In conclusion, our results suggest that, in contrast to PRORP and ELAC2 that both evolved a dependence on TRMT10C-SDR5C1 to recognize divergent mt-tRNAs, TRNT1 evolved a relaxed recognition mode to compensate for the mutational erosion of mt-tRNAs and allow a single CCA-adding enzyme to accommodate the complete sequence and structural diversity of canonical and non-canonical nu- and mt-tRNAs in the human cell.

In conjunction with previous work,^39,40,43–46,73^ our results provide a complete picture of the mitochondrial tRNA maturation pathway, in which 5’-processing by PRORP, 3’-processing by ELAC2 and 3’CCA addition by TRNT1 are sequentially catalyzed on TRMT10C-SDR5C1-bound pre-mt-tRNAs (Figure 7B). Interestingly, the requirement for the TRMT10C-SDR5C1 maturation platform appears to gradually decrease with each step of the maturation process. We hypothesize that this trend reflects the distinct functional requirements and evolutionary constraints imposed on each step, resulting in distinct molecular solutions to the shared problem of mutationally eroded mt-tRNAs. For the endonucleases PRORP and ELAC2, relaxed recognition of divergent mt-tRNAs would increase the risk of off-target cleavage in tRNA-like elements. The dependence on TRMT10C-SDR5C1, which recognizes mt-tRNAs by their overall fold rather than individual sequence or structure elements,^40^ thus provides a necessary checkpoint for processing enzymes to ensure that cleavage occurs only on mt-tRNAs as the punctuation signs between mitochondrial genes.^73,74^ By contrast, off-target 3’CCA addition by TRNT1 poses no comparable risk and occurs frequently on mitochondrial non-tRNA transcripts.^75^ Another factor may be the susceptibility of the 3’-end of mature mt-tRNAs to degradation or polyadenylation.^76–79^ The partial independence of ELAC2 and the complete independence of TRNT1 from TRMT10C-SDR5C1 may reflect the need to remove spurious poly-A tails and repair degraded 3’CCA ends to maintain a functional tRNA pool.

In summary, our results reveal the molecular basis and structural dynamics of tRNA 3’CCA addition by human TRNT1. They explain how a single CCA-adding enzyme can recognize the structurally highly divergent sets of nuclear and mitochondrial tRNAs and provide a mechanistic framework to understand the molecular etiology of disease-associated TRNT1 mutations. Finally, they show how the interplay between strong mutation pressure and the pressure to maintain an essential cellular function drives the evolution of molecular innovation.

## Methods

### Statistical methods

No statistical methods were used to predetermine sample size. The experiments were not randomized, and the investigators were not blinded to allocation during experiments and outcome assessment.

### Protein cloning, expression & purification

TRMT10C and SDR5C1 were cloned, expressed and purified as previously reported.^40^ The sequence encoding TRNT1 lacking the N-terminal mitochondrial targeting sequence (Δ1–27) was PCR amplified from human cDNA obtained from HeLa cells and cloned into the pET21b vector in frame with a C-terminal 6xHis tag. TRNT1 mutants were generated by site-directed mutagenesis using Round-the-Horn PCR amplification of wild-type TRNT1 in the pET21b vector. All TRNT1 constructs were transformed into E. coli BL21(DE3) RIL cells. Cells containing the TRNT1 plasmid were grown in LB medium at 37 °C to an OD of 0.6, followed by the induction of TRNT1 expression by the addition of IPTG to a final concentration of 0.5 mM. Cells were harvested after shaking for 16 h at 16 °C. All subsequent steps were carried out at 4 °C. Cells were harvested by centrifugation at 4000xg for 20 minutes and resuspended in lysis buffer at pH 7.6 containing 50 mM Tris/HCl, 500 mM NaCl, 30 mM imidazole, 2 mM dithiothreitol (DTT), 10% glycerol and 1x protease inhibitor cocktail (Roche). Cells were lysed by sonification, followed by centrifugation at 20,000xg for 30 minutes. The supernatant was filtered through filtration membranes with sieve width of 5 µm (Merck Millipore) and applied to a HisTrap HP 5 ml column (Cytiva) equilibrated with lysis buffer. The column was washed with 20 column volumes (CV) of lysis buffer, followed by 10 CV of low salt buffer (50 mM HEPES/NaOH, 100 mM NaCl, 30 mM imidazole, 10% glycerol, 2 mM DTT, pH 7.6). Bound protein was eluted with a gradient over 10 CV from 0-100% of elution buffer (50 mM HEPES/NaOH, 100 mM NaCl, 250 mM imidazole, 10% glycerol, 2 mM DTT, pH 7.6). Fractions containing TRNT1 according to SDS-PAGE were pooled and further purified using a Superdex 200 HiLoad 16/600 column (Cytiva) equilibrated with 20 mM HEPES/NaOH, 150 mM NaCl, 10% glycerol and 2 mM DTT at pH 7.4. Fractions containing TRNT1 were concentrated using an Amicon Ultra-4 10K Centrifugal Filter Device (Merck Millipore), and concentrated protein was aliquoted, flash-frozen in liquid nitrogen and stored at -80 °C.

The TRNT1-TRMT10C fusion construct was generated by restriction free cloning of the TRNT1(Δ1–27) sequence in frame between the TEV cleavage site and the N-terminus of TRMT10C(Δ1–38) (Figure S1A). The resulting pET14-B vector construct allowed co-expression of the TRNT1-TRMT10C fusion protein (with N-terminal 6xHis tag and TEV cleavage site) and the full-length SDR5C1. The complex of TRNT1-TRMT10C fusion and SDR5C1 was expressed and purified as described previously for the TRMT10C-SDR5C1 complex.^40^

### Preparation of substrate RNAs

All tRNA substrates were transcribed in reactions containing 1x T7 RNA polymerase reaction buffer (Thermo Fisher), 2 mM DTT, 30 mM MgCl_2_, 6 mM of each NTP, 0.002 U/l E. coli PPIase (NEB), 5 U/l T7 RNA polymerase (Thermo Fisher) and 50 ng/l template DNA. For internally FAM-labeled tRNAs used in fluorescence anisotropy experiments, the reactions were spiked with an additional 5% of FAM-UTP (Jena Bioscience). After incubation at 37°C for 10-14 hours, reactions were stopped by addition of an equal volume of 2x TBE-Urea Sample Buffer (Thermo Fisher) and boiling at 95 °C for 5 min. Samples were then separated by 12% Urea PAGE, visualized by UV-shadowing and bands containing the RNA of interest were cut and transferred to an RNase-free Eppendorf tube. Gel slices were crushed and RNAs extracted in buffer containing 300 mM NaCH_3_COO, 1 mM EDTA, and 20 mM Tris, pH 5. Extracted RNAs were ethanol-precipitated and stored at –20 °C until further use.

### 3’CCA addition assay

*In vitro* 3’CCA addition assays were carried out in buffer containing 20 mM HEPES/KOH pH 7.4, 150 mM KCl, 5 mM MgCl_2_, 0.5 mM ATP, 0.5 mM CTP, 2 mM DTT and 0.05 mg/mL BSA with 200 nM tRNA, 25 mM TRNT1, and (if applicable) 800 nM TRMT10C-SDR5C1 complex. The reactions were started by addition of TRNT1 and incubated at 30 °C for 5, 10, and 20 min. Reactions were stopped by addition of 2x TBE-Urea Sample Buffer (Thermo Fisher) supplemented with Proteinase K (Thermo Fisher). After incubation at 50 °C for 20 min and boiling for 3 min at 90 °C, samples were loaded onto a 15% TBE-Urea PAGE. Gels were subsequently soaked in TBE buffer containing 1x SYBR Gold Nucleic Acid Gel Stain (Thermo Fisher) and visualized on a Typhoon imager (Cytiva).

### Fluorescence anisotropy (FA)

FAM-labeled mt-tRNAs (20 nM) were incubated with serial dilutions of purified TRNT1, TRMT10C-SDR5C1, or TRNT1-TRMT10C-SDR5C1 complex in FA buffer (20 mM Tris/HCl pH 8, 80 mM NaCl, 40 mM KCl, 3 mM MgCl_2_, 5% glycerol, 2 mM DTT) at 20 °C for 20 min. After incubation, 20 μl of each binding reaction was transferred into a black flat-bottom 384-well microplate (Greiner) and FA measurements were performed on a Sparc Plate Reader (Tecan) using SPARKCONTROL v 3.1 with 485 nm excitation and 535 nm emission wavelengths (each with a bandwidth of 20 nm) at room temperature. Each experiment was performed in triplicates and the obtained data were analyzed using the Prism v10.2.3 (GraphPad) software. Binding curves were fitted with a single site quadratic binding equation.

### Thermal shift assay for TRNT1 variants

The thermal stability of purified TRNT1 variants was investigated by nano differential scanning fluorimetry (nanoDSF) using the Monolith NT.LabelFree Prometheus NT.48 instrument. Proteins were diluted to a concentration of 0.4 mg/ml in buffer containing 20 mM Tris/HCl pH 8, 80 mM NaCl, 40 mM KCl, 3 mM MgCl_2_ and 2 mM DTT. If applicable, ATP or CTP were added to final concentrations of 200 μM. For all experiments nanoDSF glass capillaries were used. Excitation at 280 nm (20 nm bandwidth) was set to a power of 50% yielding optimal emission intensities of 6,000-25,000 at 333-380 nm. Prior to measurement, samples were incubated at 20°C (starting temperature) for 5 min followed by heating in a thermal trap at a rate of 1°C/min from 20-95°C. The unfolding transition was monitored by changes in tryptophan and tyrosine fluorescence emission at 350/330 nm. The ratio allowed the exclusion of influences from the buffer system by showing the intrinsic changes in amino acid fluorescence. Presented melting temperatures represent the inflection points of the 350/330 nm ratio where the free energy change ΔG equals zero. All presented data result from triplicate measurements.

### Cell culture and isolation of mitochondria

Human embryonic kidney cell lines (HEK293-Flp-In T-Rex; Thermo Fisher) were cultured in high glucose (4.5 mg/ml) Dulbecco’s modified Eagle’s medium (DMEM) supplemented with 10% (v/v) fetal bovine serum (FBS) (Capricorn Scientific), 1 mM sodium pyruvate, and 2 mM L-glutamine at 37°C under a 5% CO_2_ humidified atmosphere.

For the inducible expression of C-terminally twin-strep-tagged TRMT10C, the open reading frame encoding full-length TRMT10C (NM_017819.4) was amplified using cDNA originating from HEK293 cells as template. The sequence for the twin-strep-tag was included in the reverse primer used for amplification. The resulting PCR amplicon was inserted into the pcDNA5/FRT/TO expression vector (Thermo Fisher Scientific) by restriction free cloning and the final construct was confirmed by sequencing. To generate TRMT10C-expressing cell lines, human HEK293-Flp-In T-Rex cells were transfected at ∼50% confluency with the vectors pOG44 and the pcDNA5/FRT/TO containing TRMT10C, followed by selection with Hygromycin and Blasticidin. Expression of TRMT10C^Strep^ was confirmed and optimized by titration with 0-2 g/ml tetracycline, followed by immunoblotting using TRMT10C-specific polyclonal antibody (29087-1-AP; Proteintech). To grow cells for pull-down experiments, the expression of TRMT10C^Strep^ was induced with 0.1 mg/ml tetracycline and cells were harvested after 12 h with PBS.

For mitochondrial isolation, cells were resuspended in cold TH-buffer (300 mM trehalose, 10 mM KCl, 10 mM HEPES (pH 7.4) and 0.1 mg/ml BSA) and homogenized twice. Unbroken cells were pelleted each time at 400×g for 10 min at 4°C and once at 800×g for 5 min at 4°C. The mitochondria-containing supernatant was collected and mitochondria pelleted at 10,000×g, 10 min, 4°C, pooled, once washed with BSA-free TH-buffer and the concentration determined by Bradford assay.

### Streptavidin pull-down and immunoblotting

Isolated mitochondria (1000 µg/reaction) were lysed in solubilization buffer (50 mM Tris-HCl pH 7.4, 50 mM NaCl, 10% glycerol (v/v), 1 mM EDTA, 2 mM PMSF, 1x protease inhibitor mix, 1% NP40) for 30 min at 4°C and 1000 rpm. Lysates were cleared by centrifugation (15 min, 16,000×g, 4°C) and transferred to Strep-TactinXT beads (bed 25 µl; IBA). After 1 h binding at 4°C, beads were washed 10x with washing buffer (50 mM Tris-HCl, pH 7.4, 50 mM NaCl, 10% glycerol (v/v), 1 mM PMSF, 1x PI-mix, 0.7% NP40). Bound proteins were eluted with 50 µl 1x Strep-TactinXT Elution Buffer (IBA) by shaking at 1000 rpm, 30 min at 4°C.

For immunoblotting via western blot, proteins were separated using an SDS-PAGE and afterward transferred onto PVDF membrane (Millipore) by semidry blotting. Membranes were blocked in TBST with 5% milk and then incubated with primary antibodies (TRMT10C: 29087-1-AP (Proteintech); SDR5C1: 10648-1-AP (Proteintech); ELAC2: 10071-1-AP (Proteintech); TRNT1: PA5-57878 (Thermo Fisher Scientific); VARS2: 15776-1-AP (Proteintech); SLIRP: PR3462 (homemade); MITRAC12: PR3761 (homemade); uS40m: PR5177 (homemade)) overnight at 4°C. Washed membranes were incubated with HRP-conjugated secondary antibodies (rabbit or mouse) at RT for 1 hr, washed with TBST, and visualized using the ImageQuant 800 (Cytiva) after incubation with HRP substrate.

### Cryo-EM sample preparation and data collection

All complexes were assembled in 50 μl reactions containing 1x complex preparation buffer (50 mM HEPES/KOH pH 7.0, 50 mM KCl, 0.4 mM MgCl_2_, 2 mM DTT, 100 μM SAH) with 0.4 nmol of TRNT1-TRMT10C/SDR5C1 complex (with TRNT1-TRMT10C fusion protein) and a 2-fold molar excess of mt-tRNA substrates. The complex with mt-tRNA^Tyr^ and CpCpp was assembled in the presence of 200 μM CpCpp; the complex resulting in the reconstructions with mt-tRNA^Tyr^-3’CC and -3’CCC was assembled in the presence of 200 μM CTP; the complex resulting in the reconstruction with mt-tRNA^Gln^-3’CC was assembled in the presence of 200 μM CTP and 200 μM ApCpp. All reaction mixtures were incubated at 30°C for 20 min and then applied to a Superdex 200 Increase 3.2/300 column (Cytiva) equilibrated in complex preparation buffer. Eluted fractions were analysed by SDS-PAGE and Coomassie staining. Of each diluted protein-tRNA complex (absorbance at 280 nm and a path length of 10 mm lay between 0.7-2.1 AU), 4-μl were applied to freshly glow-discharged R 2/1 holey carbon grids (Quantifoil) and the grids were blotted with a blot force of 5 for 3 s using a Vitrobot Mark IV (ThermoFisher) at 4 °C and 100% humidity immediately before plunge-freezing in liquid ethane.

Cryo-EM data collection was performed with SerialEM^80^ using a Titan Krios transmission electron microscope (Thermo Fisher Scientific) operated at 300 keV. Images were acquired in energy-filtered transmission EM mode using a GIF quantum energy filter set to a slit width of 20 eV and a K3 direct electron detector (Gatan) at a nominal magnification of x105,000, corresponding to a calibrated pixel size of 0.834 Å/pixel. Exposures were saved as non-super-resolution counting image stacks of 40 video frames, with electron doses of 0.99–1.05 e^−^/Å^2^/frame. Image stacks were acquired with stage movement per 3 × 3 holes with active beam-tilt compensation, as implemented in SerialEM.

### Cryo-EM data processing and analysis

Image stacks were preprocessed on-the-fly with gain correction, motion correction, CTF estimation, particle picking and extraction at 2.5 Å/pixel using Warp v1.0.9.^81^

For the TRNT1-mt-tRNA^Tyr^-3’•CpCpp dataset, 17,441,575 particles were autopicked from 20,780 micrographs and subjected to 2-D classification in cryoSPARC v4.7.1 (Structura Biotechnology)^82^ (Figure S2). This resulted in subsets of 8,889,056 “good” particles and 8,552,519 “bad” particles. Of the “bad” subset 200,000 particles were used for ab-initio reconstruction resulting in four “bad” references. A “good” reference was obtained from a previous smaller dataset of a TRNT1-mt-tRNA^Tyr^ complex processed in cryoSPARC. “Bad” particles were subjected to supervised 3-D classification (heterogeneous refinement algorithm in cryoSPARC) using the “good” and “bad” initial references. Of the resulting five classes, particles from three were combined and subjected to another round of 2-D classification, from which 199,303 particles were selected and combined with the initial subset of 8,889,056 “good” particles for another 2-D classification. The selected 6,690,941 particles were used to generate a “good” reference by consensus refinement for a subsequent focused 3-D classification with a mask around TRNT1 and the tRNA acceptor stem. The one class with density for TRNT1 (1,925,555 particles) was further cleaned up by heterogenous refinement using a “good” reference created by consensus refinement and “junk” references created from particle subsets by interrupted ab-initio reconstruction in which jobs were terminated shortly after the noise estimation step was completed. The resulting 1,082,468 particles were subjected to another round of focused 3-D classification with a mask around TRNT1. All particles from classes containing density for TRNT1 (three out of six) were subjected to another round of focused 3-D classification. From this, 399,714 particles were selected and re-extracted at 0.834 Å/pixel. All re-extracted particles were subjected to 2-D classification from which 24,049 “bad” particles were used to generate two “junk” references and 375,665 selected particles were used to generate a “good” reference by ab-initio reconstruction. All selected particles were then subjected to heterogenous refinement using the new “junk” and “good” references, resulting in a set of 367,432 particles that were further used for a non-uniform refinement, followed by a local refinement with a mask around TRNT1. To classify with respect to heterogeneity around the TRNT1-binding site, focused 3-D classification was carried out the same mask around TRTN1, resulting in three classes containing 121,236 (33%), 120,352 (33%) and 125,844 (34%) particles, respectively, each with robust density for TRNT1. In each of the three classes TRNT1 is bound in a slightly distinct orientation but in the same register relative to the tRNA (i.e., no translocation). Particles from the third class (125,844 particles) were further subjected to a non-uniform refinement, CTF refinement and homogenous refinement. Finally, the particles were subjected to focused refinements centered either on the TRMT10C-SDR5C1 or the TRNT1 density. The resulting TRMT10C-SDR5C1-and TRNT1-focused maps were combined using PHENIX (v1.20.1-4487),^83^ resulting in a composite map of the TRNT1 complex with mt-tRNA^Tyr^-3’•CpCpp and TRMT10C-SDR5C1.

For the TRNT1-mt-tRNA^Tyr^-3’CC/-3’CCC dataset, 31,271,743 particles were autopicked from 34,803 micrographs and subjected to 2-D classification in cryoSPARC v4.7.1^82^ (Figure S3). This resulted in a subset of 10,264,571 “good” particles and 21,007,172 “bad” particles. Of the “bad” subset 200,000 particles were used for ab-initio reconstruction resulting in four “bad” references. A “good” reference was obtained from a previous smaller dataset of a TRNT1-mt-tRNA^Tyr^ complex processed in cryoSPARC. All 10,264,571 “good” particles were subjected to heterogeneous refinement using two “good” and four “bad” initial references. Of the two resulting “good” classes, the one apparently lacking density for TRNT1 was subjected to another round of 2-D classification and the resulting “good” subset of 3,958,838 particles were combined with the second “good” class (with density for TRNT1) for another heterogenous refinement with one “good” and two “bad” references. The resulting single “good” class containing 7,491,682 particles was subjected to a supervised 3-D classification focused on the region around the TRNT1, resulting in one class with and one class without density for TRNT1. The 4,067,392 particles belonging to the class with TRNT1 density were re-extracted at 0.834 Å/pixel and subsequently subjected to 2-D classification. The resulting 4,006,063 selected particles were used to generate a “good” reference by consensus refinement, which was then used for clean-up in four consecutive rounds of heterogenous refinement, giving a final set of 3,089,164 particles. To classify with respect to heterogeneity around TRNT1, focused 3-D classification was carried out with a mask around TRNT1 and the tRNA acceptor stem, resulting in three classes containing 1,267,991 particles (class 1), 1,112,234 particles (class 2), and 708,939 particles (class 3), respectively. Class 1 and classes 2+3 contained particles with TRNT1 bound to mt-tRNA^Tyr^ in different states, with TRNT1 translocated by one register along the acceptors stem in classes 2+3 relative to class 1. Each of the three classes was subjected to another round of focused 3-D classification with masks around TRNT1 and the acceptor stem. Each gave rise to three “sub-classes” with TRNT1 bound in slightly distinct orientations but retaining the same register relative to the tRNA (i.e., no translocation). First, a single sub-class of class 1 containing 373,380 particles was further subjected to a non-uniform refinement and orientation rebalancing, giving a final set of 127,197 particles. These particles subjected to CTF refinement and homogenous refinement and a focused refinement centered on the TRNT1 density. The resulting consensus and TRNT1-focused maps were combined using PHENIX,^83^ resulting in a composite map of the TRNT1 complex with mt-tRNA^Tyr^-3’CC and TRMT10C-SDR5C1. Second, one sub-class of class 2 containing 398,366 particles and one sub-class of class 3 containing 213,457 particles were combined and subjected to the same procedure as described above, resulting in a composite map of the TRNT1 complex with mt-tRNA^Tyr^-3’CCC and TRMT10C-SDR5C1 from a final 340,521 particles.

For the TRNT1-mt-tRNA^Gln^-3’CC dataset, 19,293,885 particles were autopicked from 23,439 micrographs and subjected to 2-D classification in cryoSPARC v4.7.1^82^ (Figure S4). This resulted in a subset of 8,085,967 “good” particles and 11,207,918 “bad” particles. Of the “bad” subset 200,000 particles were used for ab-initio reconstruction resulting in four “bad” references. A “good” reference was obtained from a previous smaller dataset of a TRNT1-mt-tRNA^Tyr^ complex processed in cryoSPARC. All 8,085,967 “good” particles were subjected to supervised heterogeneous refinement using two “good” and four “bad” initial references. 4,485,418 particles belonging to two “good” classes were subjected to a supervised 3-D classification focused on the region around the TRNT1, followed by three rounds of heterogenous refinement with “junk” classes generated by interrupted ab-initio reconstruction. This resulted in a set of 1,016,167 particles that were re-extracted at 0.834 Å/pixel. A subset of the re-extracted particles was subjected to 2-D classification from which 26,871 “bad” particles were used to generate “junk” references by ab-initio reconstruction and 116,782 “good” particles were used to generate a “good” reference by consensus refinement. All 1,016,167 re-extracted particles were then subjected to another round of heterogenous refinement using the new “junk” and “good” references, resulting in a set of 882,441 particles that were used for a subsequent non-uniform refinement. To classify with respect to heterogeneity around the TRNT1-binding site, focused 3-D classification was carried out with a mask around TRTN1 and the tRNA acceptor stem. Focused 3-D classification with the same mask was repeated with 629,579 particles, resulting in three classes containing 193,434 (34%), 185,219 (33%) and 190,276 (33%) particles, respectively, each with robust density for TRNT1. In each of the three classes TRNT1 is bound in a slightly distinct orientation but in the same register relative to the tRNA (i.e., no translocation). Particles from the third class (190,276 particles) were further subjected to a non-uniform consensus refinement, CTF refinement and homogenous refinement. Due to strong orientation bias in the particle set, orientations were rebalanced, resulting in a final set of 99,883 particles. These particles were subjected to a consensus 3-D refinement and a focused refinement centered on the TRNT1 density. The resulting consensus and TRNT1-focused maps were combined using PHENIX,^83^ resulting in a composite map of the TRNT1 complex with mt-tRNA^Gln^-3’CC and TRMT10C-SDR5C1.

For all reconstructions, local resolution estimations and angular distribution plots were calculated in cryoSPARC v4.7.1.^82^

### Model building and refinement

The initial models for TRMT10C, SDR5C1, mt-tRNA^Tyr^, and mt-tRNA^Gln^ were obtained from the models of the mt-RNase Z complex (PDBs: 8rr1 and 8rr3),^39^ and for TRNT1 from AlphaFoldDB (AF-Q96Q11). The models were rigid-body-fit into the final maps of the complexes of TRNT1 bound to mt-tRNAs and TRMT10C-SDR5C1 using UCSF ChimeraX 1.7.1 and rebuilt and refined using Coot v0.9.8.92.^84^ The complete models of mitochondrial CCA-adding enzyme complexes were refined against the respective composite maps using phenix.real-space refine in PHENIX (v1.20.1-4487).^83^ The MolProbity package within PHENIX suite was used for model validation.^85^ All structural analyses and image renderings for figure preparation were done using UCSF ChimeraX 1.7.1.^86^

## Data availability

The uniprot accession IDs for TRNT1, TRMT10C and SDR5C1 are Q96Q11, Q7L0Y3 and Q99714. The structure coordinates for mt-RNase P and mt-RNase Z were obtained from Protein Data Bank (PDB) under accession codes 7onu, 8rr1, 8rr3, and 8rr4. The cryo-EM density reconstructions for the mitochondrial CCA-adding enzyme complexes with mt-tRNA^Gln^-3’CC, mt-tRNA^Tyr^-3’A_73_ and CpCpp, mt-tRNA^Tyr^-3’CC, and mt-tRNA^Tyr^-3’CCC were deposited with the Electron Microscopy Database (EMDB) under accession codes EMD-56474, EMD-56492, EMD-56502, and EMD-56514, respectively. The respective structure coordinates were deposited with the Protein Data Bank (PDB) under accession codes 9TZQ, 9U0I, 9U0S, and 9U14. Source data are provided with this manuscript.

## Acknowledgements

We thank all members of the Hillen Lab for discussion, Christian Dienemann and Ulrich Steuerwald (MPI-NAT cryo-EM facility) for assistance with cryo-EM data acquisition, and Johannes Sattmann for assistance with pull-down experiments. This work was supported by the Deutsche Forschungsgemeinschaft under Germany’s Excellence Strategy EXC 2067/1-390729940, FOR2848, SFB1565 (project number 469281184, P13 and P14) (to H.S.H. and P.R.) and the priority program SPP2453 (project number 541758684) (to S.D.), by the European Union (ERC Starting Grant MitoRNA, grant agreement no. 101116869 to H.S.H.; ERC Advanced Grants MiXpress, grant agreement no. 101054637 to P.R.), and by the EMBO Young Investigator Programme (to H.S.H.). Views and opinions expressed are however those of the author(s) only and do not necessarily reflect those of the European Union or the European Research Council Executive Agency. Neither the European Union nor the granting authority can be held responsible for them.

## Author contributions

B.K. carried out all experiments unless stated otherwise. L.K. purified mutant TRNT1 proteins and performed thermal shift assays. A.B. assisted in single-particle cryo-EM analysis. S.D. carried out immunoprecipitation experiments. B.K., P.R. and H.S.H. conceptualized research. P.R. and H.S.H. supervised research. B.K., S.D. and H.S.H interpreted the data. B.K. and H.S.H wrote the manuscript.

## Supplemental Figures and Tables

**Supplemental Figure S1:**
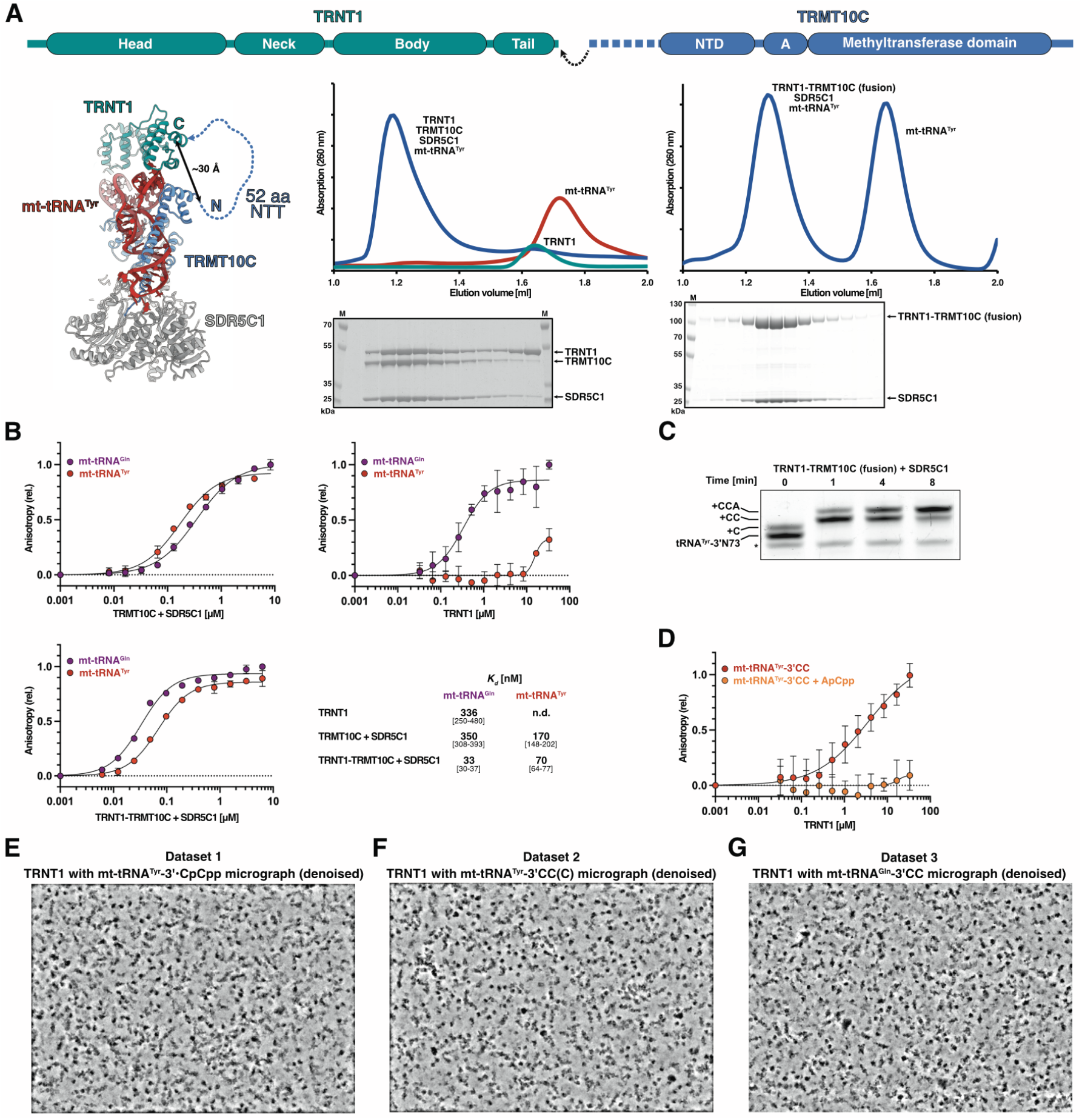
*In vitro* reconstitution of mitochondrial CCA-adding complexes, related to Figure 1. **(A)** Design of the TRNT1-TRMT10C fusion protein. The N-terminus of TRMT10C(Δ1–38) was fused directly to the C-terminus of TRNT1(Δ1–27). In the structural model of TRNT1 bound to TRMT10C-SDR5C1 and mt-tRNA^Tyr^ (left; PDB: 9u0i), the first ordered residue of the TRMT10C NTD is Ala_92_, which lies ∼30 Å from the C-terminus of TRNT1 (black arrow). The N-terminal 52 residues of TRMT10C (aa 39-91; dashed blue line) are disordered and not resolved in the cryo-EM map. Chromatograms show the SEC elution profiles for mitochondrial CCA-adding complexes (blue) reconstituted with mt-tRNA^Tyr^ and either individual proteins (middle) or the TRNT1-TRMT10C fusion protein (right). The samples loaded onto the SDS-PAGEs (bottom) are aligned with the corresponding elution volumes in the chromatograms above. **(B)** Fluorescence anisotropy of FAM-labelled mt-tRNA^Gln^ and mt-tRNA^Tyr^ precursors with TRNT1, TRMT10C/SDR5C1 and the TRNT1-TRMT10C/ SDR5C1 fusion complex. Each dot represents the mean from three replicates with standard deviations shown as error bars. Protein-RNA dissociation constants (K_d_) were determined by fitting the data with a single site quadratic binding equation (Methods). The binding data did not provide a reliable estimate for the K_d_ between mt-tRNA^Tyr^ and TRNT1. For each determined K_d_ value, the corresponding 95% confidence intervals are given in brackets. **(C)** *In vitro* CCA-addition assay with the TRNT1-TRMT10C/SDR5C1 fusion complex and mt-tRNA^Tyr^. The asterisk marks an RNA band corresponding to the length of mt-tRNA^Tyr^ lacking the terminal 3’A_73_ nucleotide. **(D)** Fluorescence anisotropy of FAM-labelled mt-tRNA^Tyr^-3’CC (lacking only the terminal A_76_) with TRNT1 either in the absence (red) or presence (orange) of 200 M ApCpp. Each dot represents the mean from three replicates with standard deviations shown as error bars. For improved binding of TRNT1 to mt-tRNA^Tyr^, the salt concentration was reduced compared to standard FA buffer to 30 mM NaCl and 20 mM KCl. **(E-G)** Denoised representative cryo-EM micrographs of the CCA-adding complex containing CpCpp and mt-tRNA^Tyr^-3’ (state 1) from a total of 20,780 collected micrographs (E), of the complex containing mt-tRNA^Tyr^-3’CC (state 2) or mt-tRNA^Tyr^-3’CCC (state 3) from a total of 34,803 collected micrographs (F), and of the complex containing mt-tRNA^Gln^-3’CC (state 2’) from a total of 23,439 collected micrographs (G),

**Supplemental Figure S2:**
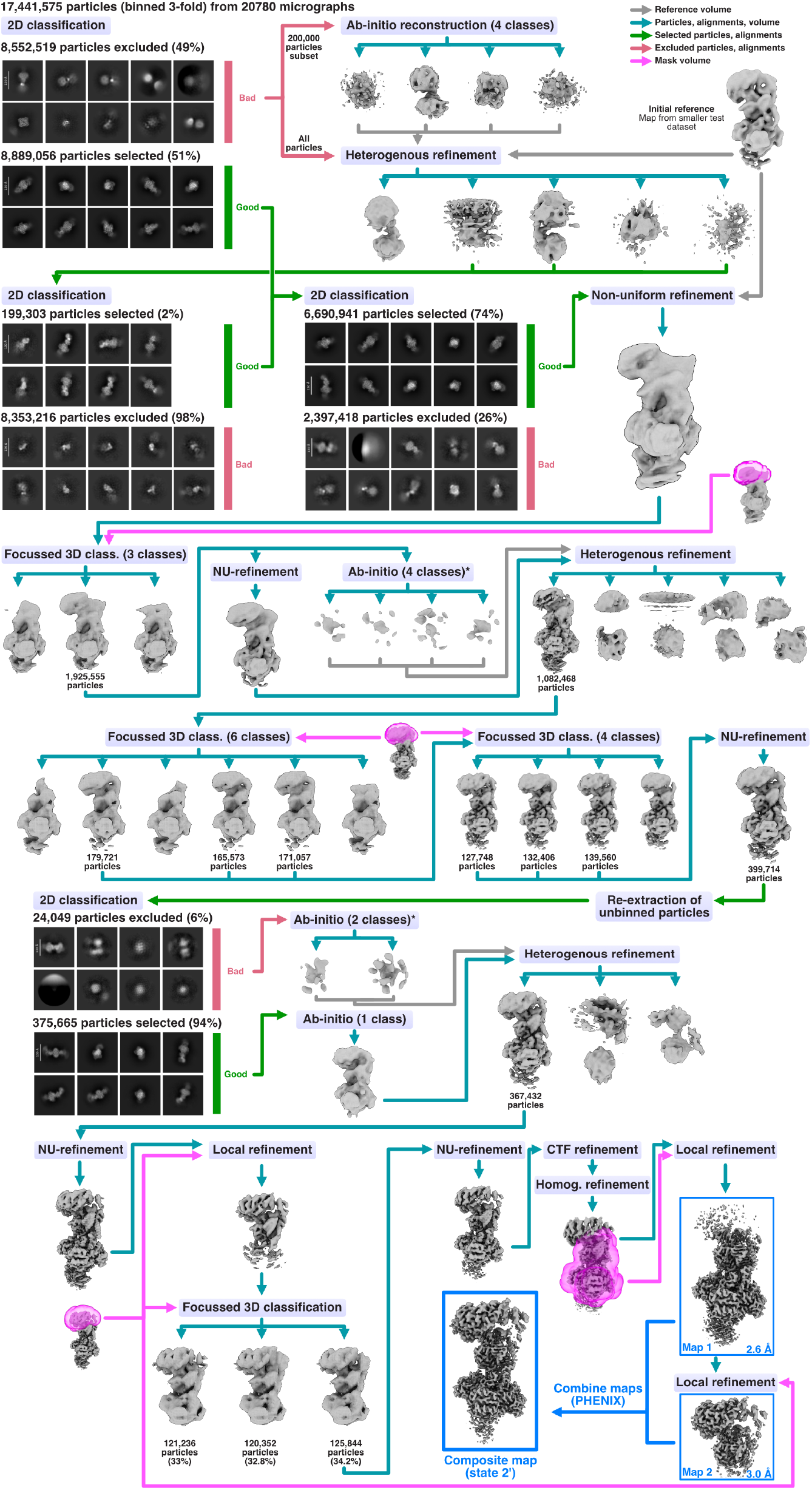
Cryo-EM processing workflow for the mitochondrial CCA-adding complex containing mt-tRNA^Tyr^-3’ and CpCpp (state 1), related to Figure 1. * indicates ab-initio reconstruction jobs that were terminated shortly after the noise estimation step was completed.

**Supplemental Figure S3:**
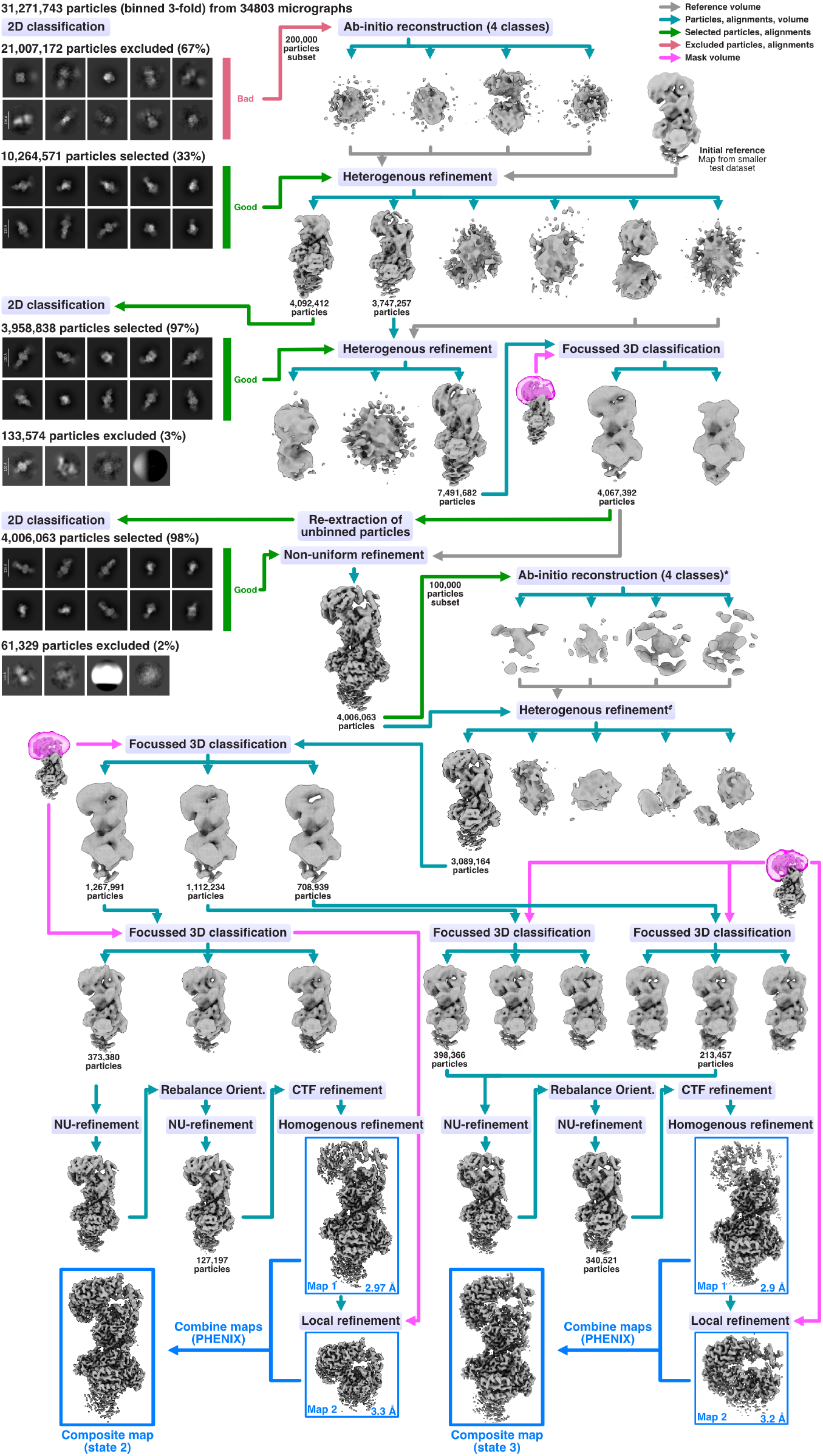
Cryo-EM processing workflow for the mitochondrial CCA-adding complexes containing mt-tRNA^Tyr^-3’CC and mt-tRNA^Tyr^-3’CCC (states 2 and 3), related to Figure 1. * indicates ab-initio reconstruction jobs that were terminated shortly after the noise estimation step was completed. # marks heterogenous refinement jobs that were repeated four times to remove junk particles; shown are only the volumes from the final refinement job.

**Supplemental Figure S4:**
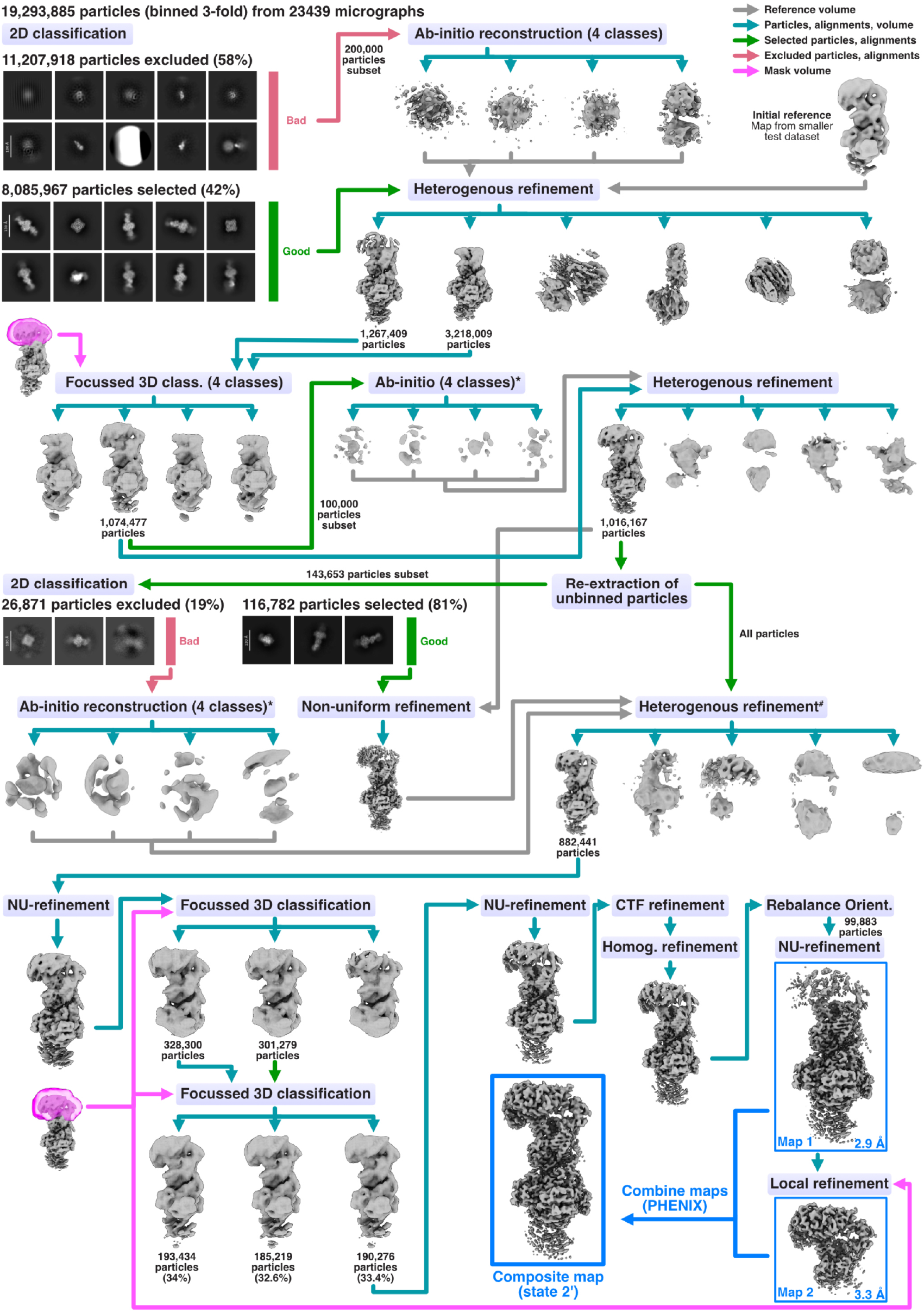
Cryo-EM processing workflow for the mitochondrial CCA-adding complex containing mt-tRNA^Gln^-3’CC (state 2’), related to Figure 1. * indicates ab-initio reconstruction jobs that were terminated shortly after the noise estimation step was completed. # marks heterogenous refinement jobs that were repeated three times to remove junk particles; shown are only the volumes from the final refinement job.

**Supplemental Figure S5:**
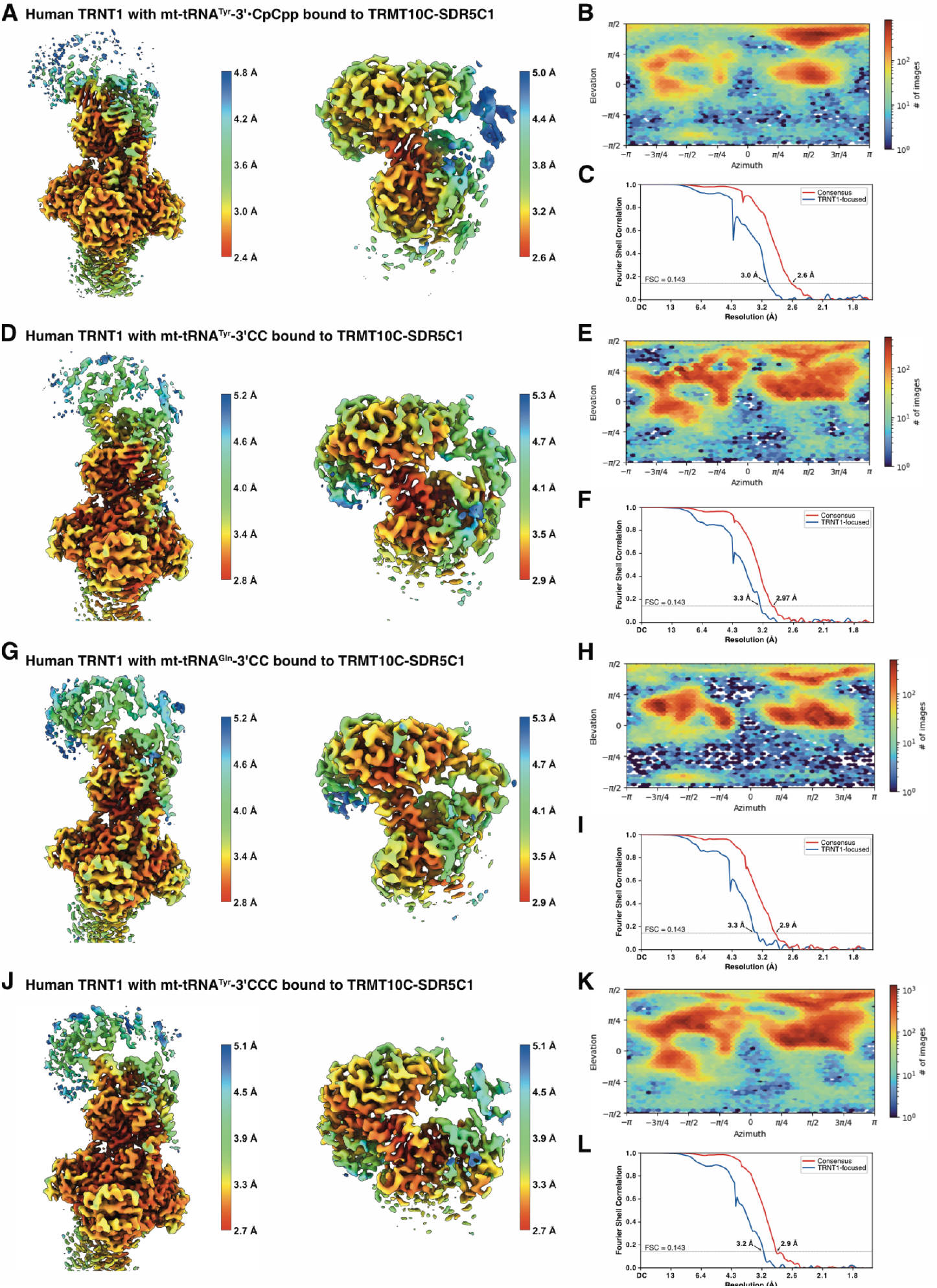
Cryo-EM post-processing analysis, related to Figure 1. **(A)** Local resolution maps for the TRMT10C-SDR5C1-focused map (left) and TRNT1-focused map (right) of the mitochondrial CCA-adding complex containing mt-tRNA^Tyr^-3’ and CpCpp (state 1). **(B)** Angular distribution plots for the TRMT10C-SDR5C1-focused map in (A). **(C)** Corrected Fourier Shell Correlation (FSC) plots for the maps in (A). FSC threshold of 0.143 was used for reporting resolutions. **(D)** Local resolution maps for the consensus refinement map (left) and TRNT1-focused map (right) of the mitochondrial CCA-adding complex containing mt-tRNA^Tyr^-3’CC (state 2). **(E)** Angular distribution plots for the consensus map in (D). **(F)** Corrected Fourier Shell Correlation (FSC) plots for the maps in (D). **(G)** Local resolution maps for the consensus refinement map (left) and TRNT1-focused map (right) of the mitochondrial CCA-adding complex containing mt-tRNA^Gln^-3’CC (state 2’). **(H)** Angular distribution plots for the consensus map in (G). **(I)** Corrected Fourier Shell Correlation (FSC) plots for the maps in (G). **(J)** Local resolution maps for the consensus refinement map (left) and TRNT1-focused map (right) of the mitochondrial CCA-adding complex containing mt-tRNA^Tyr^-3’CCC. **(K)** Angular distribution plots for the consensus map in (J). **(L)** Corrected Fourier Shell Correlation (FSC) plots for the maps in (J).

**Supplemental Figure S6:**
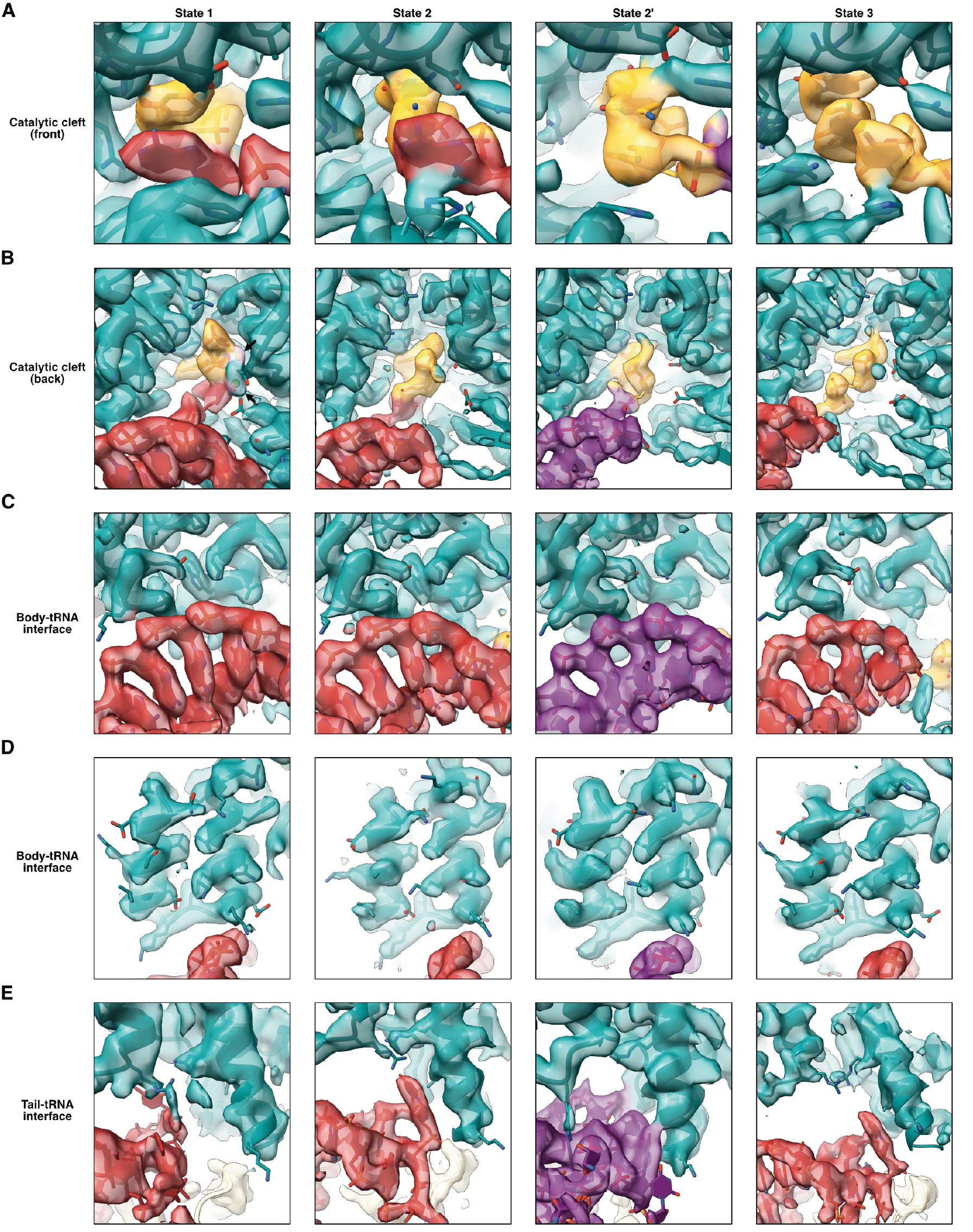
Details of cryo-EM models and densities, related to Figures 1 and 2. **(A)** Cryo-EM models and densities around the entrance to the catalytic cleft of TRNT1. The colouring scheme is the same as in Figure 1B. **(B)** Cryo-EM models and densities in the catalytic cleft and active site pocket of TRNT1. Arrows in state 1 indicate the additional density between the phosphate moieties of the CpCpp molecule and the active site residues of TRNT1 interpreted as Mg^2+^ ions. **(C)** Cryo-EM models and densities for the interface between the TRNT1 body domain and the acceptor stem of the bound mt-tRNA. **(D)** Cryo-EM models and densities for helices 14 and 15 of the TRNT1 body domain and the acceptor stem of the bound mt-tRNA. **(E)** Cryo-EM models and densities for the interface between the TRNT1 tail domain and the elbow region of the bound mt-tRNA.

**Supplemental Figure S7:**
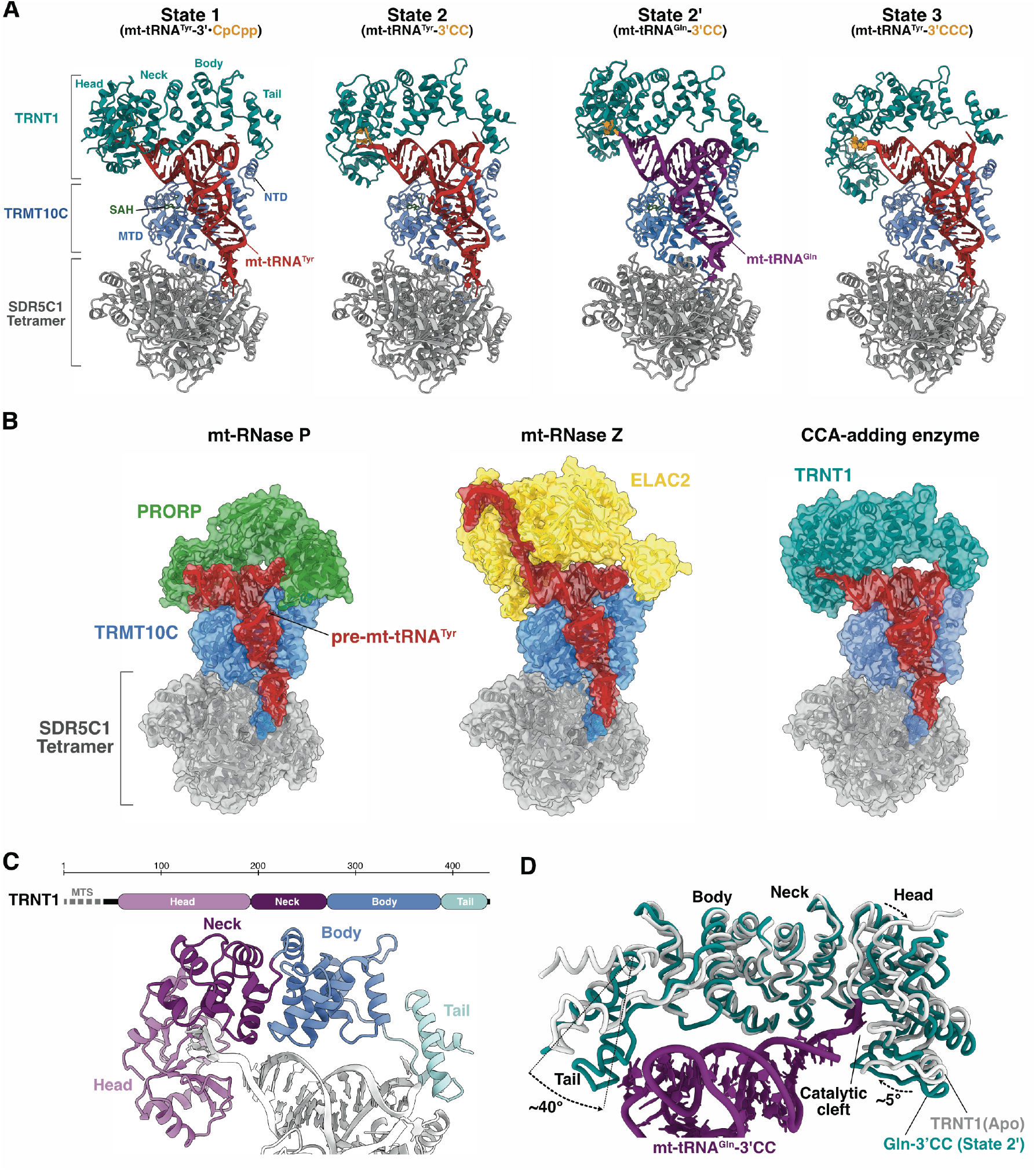
Structures of mitochondrial CCA-adding enzyme complexes, related to Figures 1 and 2. **(A)** Overall structural models for the four reconstructed states of the mitochondrial CCA-adding enzyme complex. The coloring scheme is the same as in Figure 1. SAH in the active site of TRMT10C in all four states is shown as green sticks. **(B)** Comparison of the overall structures of the human mitochondrial RNase P complex with PRORP (green; PDB: 7onu),^1^ the RNase Z complex with ELAC2 (yellow; PDB: 8rr4),^2^ and the CCA-adding complex with TRNT1 (teal) bound to mt-tRNA^Tyr^. **(C)** Domain architecture of TRNT1. The tRNA is shown in white. **(D)** Conformational rearrangement (indicated by dashed arrows) in the head and tail domains of TRNT1 (teal) bound to mt-tRNA^Gln^-3’CC (state 2’) relative to the crystal structure of apo TRNT1 (white; PDB: 4×4w).^3^

**Supplemental Figure S8:**
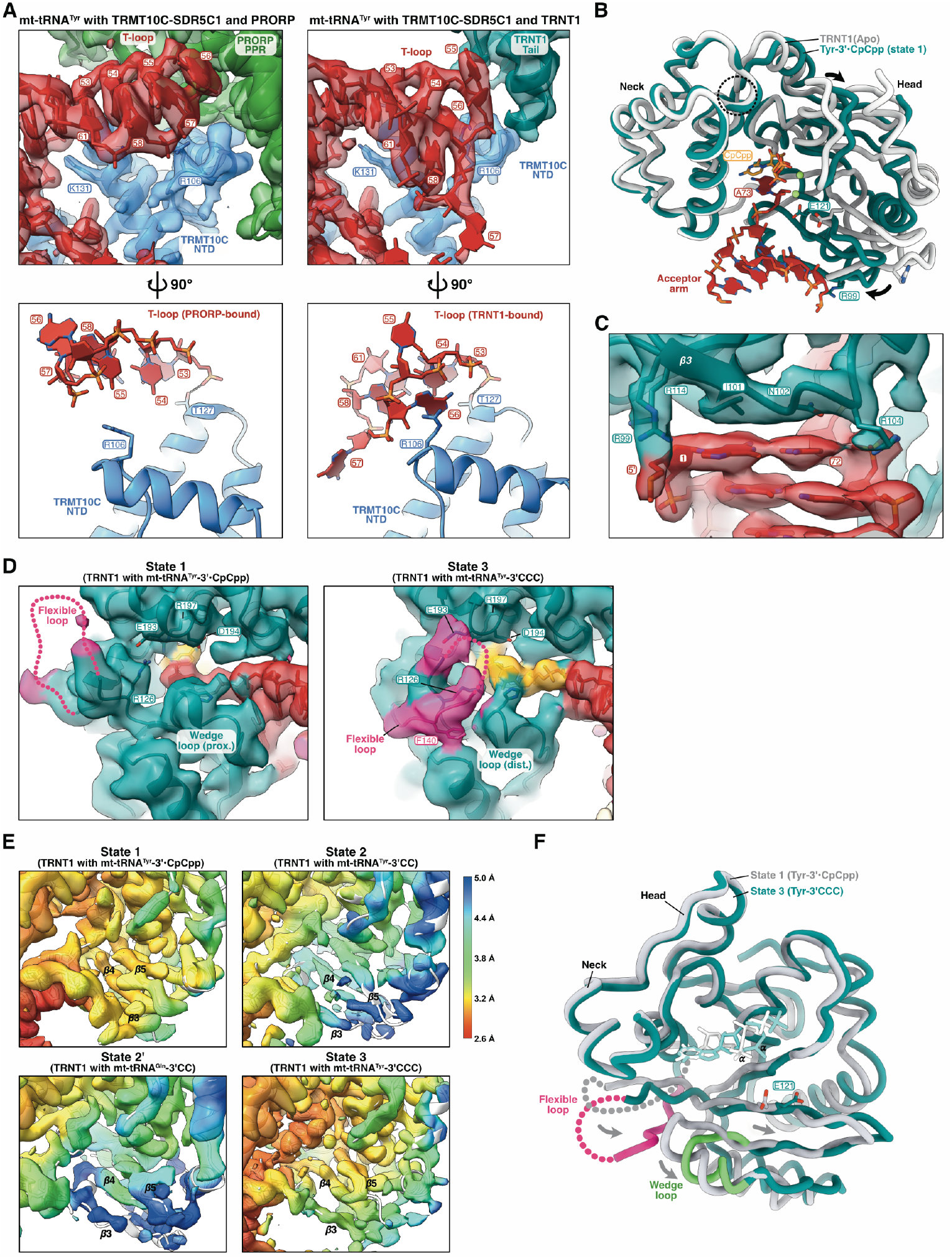
Details of mitochondrial CCA-adding enzyme structures, related to Figure 3. **(A)** The T-loop of mt-tRNA^Tyr^ (red) adopts distinct structures in the mitochondrial RNase P (left; PDB: 7onu; EMDB: 13002)^1^ and CCA-adding complexes (right). In the lower panels the hydrogen bond of Thr_127_ to the phosphate moiety of G_53_ is shown for reference. **(B)** Substrate-induced closure of the head domain relative to the neck domain in mt-tRNA^Tyr^-3’•CpCpp bound TRNT1 (teal) relative to apo TRNT1 (white; PDB: 4×4w).^3^ The superposition is based on the neck domain. The hinge region for the closure movement of the head is indicated by a dashed circle. In both structures Arg_99_ at the *N*-terminus of β_3_ and the catalytic Glu_121_ in β_5_ are shown as sticks for reference. **(C)** Model and cryo-EM map for the interface between the terminal base-pair of the tRNA acceptor stem (G1:C72) and strand 3 of the TRNT1 head domain in state 1. **(D)** Model and cryo-EM map (unsharpened) for the entrance to the catalytic cleft in state 1 (left) and state 3 (right), with the model and density for the flexible loop region colored in pink. **(E)** Comparison of the local resolution maps (TRNT1-focused map) for the head domain of TRNT1 in the four reconstructed states. **(F)** Comparison of the structural arrangements in the head domain between state 1 (white) and state 3 (teal). CpCpp and ATP (modeled as in Figure 3F) are shown as white and cyan sticks, respectively.

**Supplemental Figure S9:**
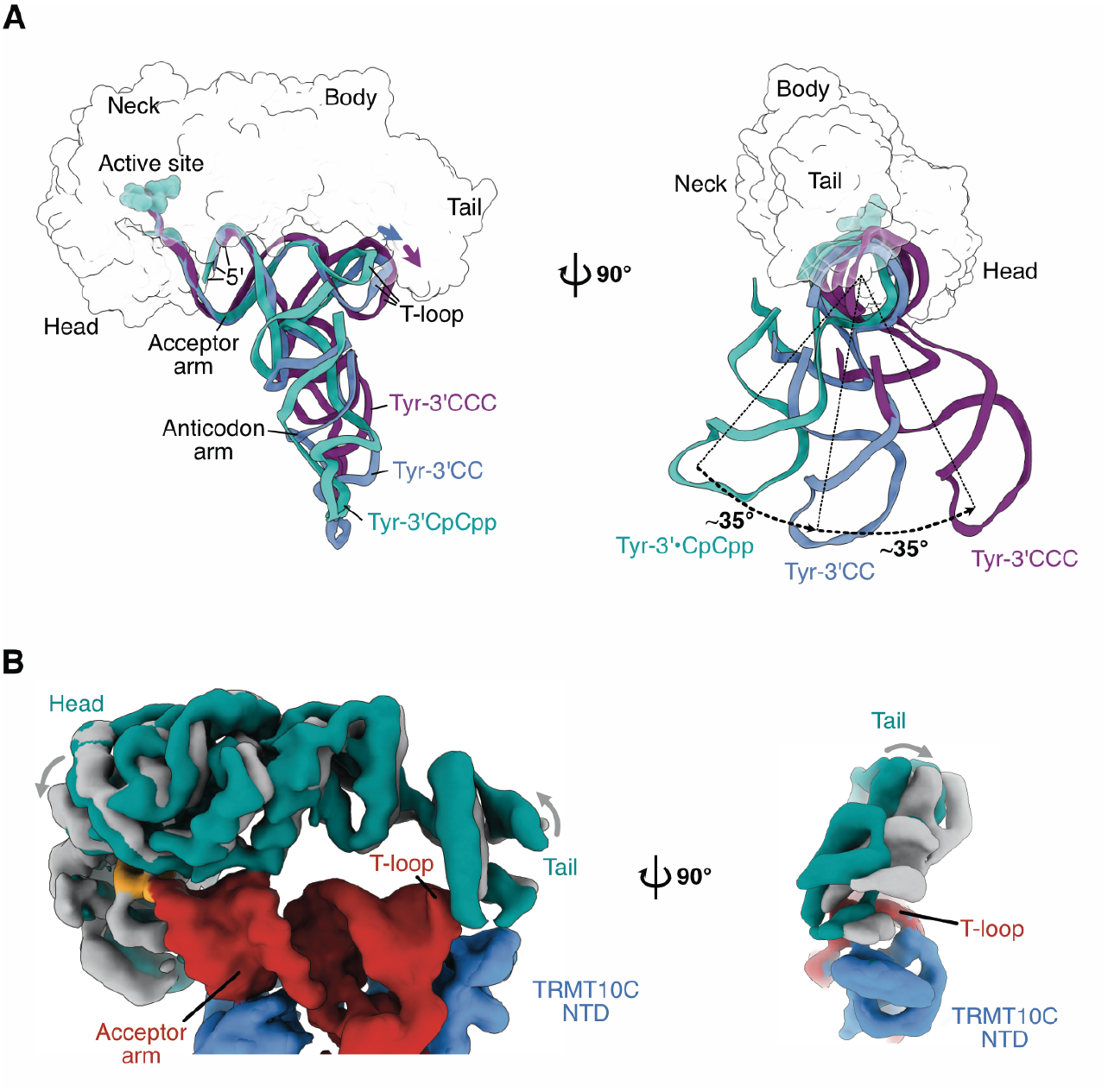
Translocation of TRNT1 and tRNA during 3’-extension, related to Figure 4. **(A)** Translocation of the tRNA relative to TRNT1. Structures are superimposed on TRNT1 (shown as transparent surface). Mt-tRNA^Tyr^ is colored cyan (state 1), blue (state 2), and purple (state 3), respectively. Arrows in the top panel indicate the ‘sliding’ movement of the tRNA’s T-loop relative to the tail domain between states 1 and 2 (blue) and states 2 and 3 (purple). **(B)** Maps (unsharpened) of additional sub-states of state 3, containing mt-tRNA^Tyr^-3’CCC, showing that further rotation of TRNT1 relative to the tRNA minihelix domain (acceptor-arm and T-arm) results in the gradual loss of interactions between tail domain and T-loop. The rotation of TRNT1 between these sub-states does not involve a translocation of TRNT1 relative to the tRNA.

**Supplemental Figure S10:**
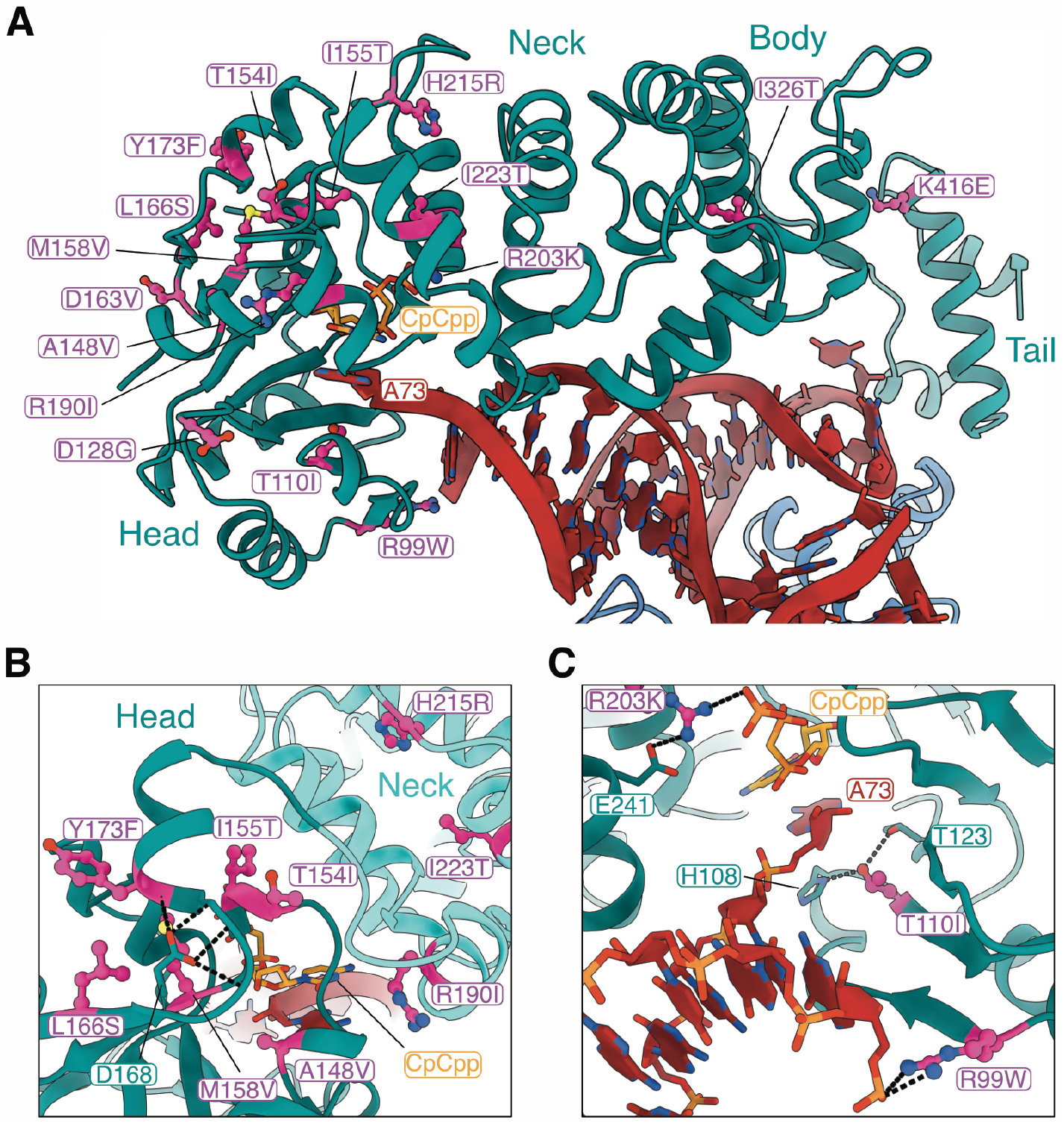
Disease-causing TRNT1 mutations, related to Figure 6. **(A)** Known disease-causing mutations^4^ mapped onto the model of TRNT1 in state 1. Mutation sites are highlighted in pink with wild-type side chains shown in ball-and-stick presentation. Nonsense mutations are not included. **(B)** Close-up of disease-causing mutation sites in the interface of head (teal) and neck (cyan) domains. Most mutations cluster in regions that are likely involved in interdomain dynamics during 3’CCA addition. Mutation of Asp_168_ (teal sticks) in the same region was previously shown to affect interdomain movements and interfere specifically with A_76_ addition^5^. **(C)** Close-up of disease-causing mutation sites in the catalytic cleft of TRNT1 that likely affect substrate binding and/or the nucleotidyl-transfer reaction directly.

**Supplemental Table 1.** Cryo-EM data collection, refinement and validation statistics.

|  | Dataset 1 | Dataset 2 |  | Dataset 3 |
| --- | --- | --- | --- | --- |
|  | State 1 | State 2 | State 3 | State 2' |
|  | EMDB-56492 | EMDB-56502 | EMDB-56514 | EMDB-56474 |
|  | PDB 9u0i | PDB 9u0s | PDB 9u14 | PDB 9tzq |
| **Data collection and processing** |  |  |  |  |
| Magnification | x105,000 | x105,000 |  | x105,000 |
| Acceleration voltage (kV) | 300 | 300 |  | 300 |
| Electron exposure (e<sup>-</sup>/Å<sup>2</sup>) | 50.00 | 50.01 |  | 49.99 |
| Defocus range (μm) | 0.5 – 2.5 | 0.5 – 2.5 |  | 0.5 – 2.5 |
| Pixel size (Å) | 0.834 | 0.834 |  | 0.834 |
| Symmetry imposed | C1 | C1 |  | C1 |
| Initial particle images (no.) | 17,441,575 | 31,271,743 |  | 19,293,885 |
| Final particle images (no.) | 125,844 | 127,197 | 340,521 | 99,883 |
| Map resolution (Å) | 2.6 (Map 1) | 3.0 (Map 1) | 2.9 (Map 1) | 2.9 (Map 1) |
|  | 3.0 (Map 2) | 3.3 (Map 2) | 3.2 (Map 2) | 3.3 (Map 2) |
| FSC threshold | 0.143 | 0.143 | 0.143 | 0.143 |
| Map sharpening *B* factors (Å<sup>2</sup>) | -44.6 (Map 1) | -74.9 (Map 1) | -87.5 (Map 1) | -79.8 (Map 1) |
|  | -83.0 (Map 2) | -112.6 (Map 2) | -127.6 (Map 2) | -120.4 (Map 2) |
| **Refinement** |  |  |  |  |
| Initial models | State 2 | PDB: 8rr4<br>AF-Q96Q11 | State 2 | PDB: 8rr3<br>AF-Q96Q11 |
| Model resolution (Å) | 2.9 | 3.2 | 3.0 | 3.1 |
| FSC threshold | 0.5 | 0.5 | 0.5 | 0.5 |
| CC (masked) | 0.89 | 0.90 | 0.89 | 0.88 |
| **Model composition** |  |  |  |  |
| Non-hydrogen atoms | 14,116 | 14,251 | 14,337 | 13,952 |
| Protein residues | 1,668 | 1,685 | 1,693 | 1,632 |
| Nucleotide residues | 64 | 66 | 67 | 71 |
| Ligands | 2TM: 1<br>MG:2<br>SAH:1 | SAH:1 | SAH:1 | SAH:1 |
| **Mean *B* factors (Å<sup>2</sup>)** |  |  |  |  |
| Protein | 42.18 | 66.07 | 57.34 | 100.18 |
| Nucleotide | 77.78 | 98.86 | 107.07 | 137.35 |
| Ligand | 35.44 | 41.23 | 28.16 | 53.14 |
| **RMS deviations** |  |  |  |  |
| Bond lengths (Å) | 0.004 | 0.002 | 0.003 | 0.003 |
| Bond angles (°) | 0.656 | 0.559 | 0.536 | 0.628 |
| **Validation** |  |  |  |  |
| MolProbity score | 1.45 | 1.43 | 1.40 | 1.53 |
| Clash score | 6.37 | 7.88 | 6.02 | 9.13 |
| Rotamer outliers (%) | 0.82 | 0.07 | 0.37 | 0.0 |
| **Ramachandran plot** |  |  |  |  |
| Favored (%) | 97.51 | 98.44 | 97.67 | 97.82 |
| Allowed (%) | 2.49 | 1.56 | 2.33 | 2.18 |
| Outliers (%) | 0.0 | 0.0 | 0.0 | 0.0 |

